# Phylogenomic analyses of the diverse desert-alpine plant lineage Cistantheae

**DOI:** 10.1101/2025.10.10.680062

**Authors:** Anri Chomentowska, Sophie G. Dauerman, Nora Heaphy, Leonardo Gaspar, Pablo M. Molina, C. Matt Guilliams, R. Matthew Ogburn, Lillian P. Hancock, Joseph AM Holtum, Andrés Moreira-Muñoz, Mónica Arakaki, Patrick W. Sweeney, Iris E. Peralta, Erika J. Edwards

## Abstract

Deserts and alpine habitats, though ecologically distinct, share similar environmental stressors such as drought and high radiation. Various plant lineages traverse both biomes, which is often associated with transitions in life history strategy, where annuality is more often associated with drier desert habitats and perenniality more common in higher elevations. One such lineage is Cistantheae (Montiaceae), a morphologically diverse herbaceous clade in western North and South America. We aimed to infer a robust phylogeny of the clade as a foundation for taxonomic and comparative work. We used double-digest RADSeq to generate reduced-representation genomic data from over 160 samples representing 48 putative species in Cistantheae. Maximum likelihood and coalescent-based phylogenetic methods were utilized to infer evolutionary relationships across the full clade and within major subclades. We tested for signatures of admixture and introgressive gene flow, and reconstructed ancestral life history and climate niche to identify patterns of correlated evolution. We inferred a well-resolved phylogeny of Cistantheae, providing strong support for relationships among subclades within Cistantheae. While many species relationships were clarified, we also found evidence of rampant gene flow and incomplete lineage sorting, particularly within the annual *Cistanthe* clade from the Atacama Desert. Life history is evolutionarily labile across the clade, and was strongly correlated with temperature/ precipitation-related bioclimatic variables: annuals tend to occur in hotter, drier environments, while perennials in cooler, wetter. Elevational range was also evolutionarily labile, with several species occupying broad elevational gradients. We present the first densely-sampled, phylogenomic analysis of Cistantheae, providing key insights into species relationships in the clade. Repeated transitions in life history and climate niche, alongside wide elevational ranges, suggest that many Cistantheae species may be preadapted to both arid and montane habitats. This phylogeny will underpin further comparative, taxonomic, and phylogenomic studies in this ecologically important lineage.

## INTRODUCTION

The dramatic landscapes of western North and South America host a diversity of ecosystems in relatively close proximity to one another, including high elevation environments, hyper-arid deserts and two of the World’s five Mediterranean-type climate zones. These montane and xeric systems are noted for their high rates of endemism (Chabot and Billings 1972; Arroyo *et al*. 1988; Rundel *et al*. 1991); at the same time, botanists have also long noticed floristic connections between them (Went 1948; Sarmiento 1975). In general, the colonization of high-elevation or alpine regions and subsequent diversification were speculated to have originated from adjacent lowland or desert ancestors (Stebbins 1982; Axelrod and Raven 1985). More recently, phylogenetic studies have confirmed these connections, documenting movements of plant lineages between desert and mountain ecosystems, including *Linanthus* (Bell and Patterson 2000; Anghel *et al*. 2025)*, Cirsium* (Kelch and Baldwin 2003; Siniscalchi *et al*. 2023), *Oxalis* (Heibl and Renner 2012), and the Montiaceae (Ogburn and Edwards 2015). Although desert and alpine environments appear to be quite distinct, at least in terms of temperature (Chabot and Billings 1972), there are other factors that might make desert and alpine plants reciprocally pre-adapted; both environments present extreme radiation loads, for example, and many high elevation environments are also quite arid (Billings and Mooney 1968; Ehleringer 1985; Körner 2003). Consequently, there are overlaps in physiological effects of both cold and drought stress (e.g., negative turgor and change in membrane potential at the cellular level; Beck *et al*. 2007) and the plants’ responses to these stressors (e.g., stomatal closure; Agurla *et al*. 2018).

Multiple studies that documented plant transitions between desert and alpine systems also noted that these shifts are often correlated with changes in life history, with annuals occupying lowland arid areas and perennials more common in the sub-alpine and alpine regions. It has been long established that higher-elevation areas (associated with lower temperatures) often lack annual plant species (Körner 2003). Across 32 different angiosperm groups, temperature (specifically the highest temperature of the warmest month) influenced the evolution of annuality (Boyko *et al*. 2023). Notably, Ogburn and Edwards (2015) found that in Montiaceae, annuals tend to occupy warmer environments, and perennials tend to occupy colder. Similar associations have also been observed in other systems; in *Oenothera* (Evans *et al*. 2005), *Nemesia* (Datson *et al*. 2008), *Heliophila* (Monroe *et al*. 2019), and *Veronica* (Wang *et al*. 2016), shifts to annuality are correlated with increasing temperature, but also with increased aridity or xeric habitats. Globally, annuals have been found to be favored in dry and hot areas, and a model for biogeographical distribution showed specifically that precipitation and temperature in the driest quarter explains annual plant abundance (Poppenwimer *et al*. 2023). Conversely, though less frequent, shifts to perenniality are associated with the colonization of montane and alpine habitats in lineages such as *Lupinus* (Drummond *et al*. 2012) and Arabideae (Karl and Koch 2013).

The plant family Montiaceae (crown age ∼40 MYA; Arakaki *et al*. 2011; Caryophyllales) comprises roughly 250 species, widespread and ecologically diverse, inhabiting a range of disparate habitats in the Americas, Australia, New Zealand, and Europe. Montiaceae has been recognized as monophyletic in the process of re-circumscribing Portulacaceae *sensu lato* (Nyffeler & Eggli 2010), with phylogenetic analysis determining two main lineages in Montiaceae (Ogburn and Edwards 2015). One of these lineages includes recognized genera *Calyptridium* Nutt., *Cistanthe* Spach, *Montiopsis* Kuntze, and *Philippiamra* Kuntze (*sensu* Hershkovitz 2019; previously *Cistanthe* sect. *Philippiamra*; Hershkovitz 1990), and monotypic *Lenzia* Phil and *Thingia* Hershk (Hershkovitz 2019), comprising the Cistantheae clade (Hershkovitz 2019). Within Cistantheae, *Montiopsis*, *Philippiamra*, and *Lenzia* are exclusively South American clades––distributed across Chile, Peru, western Argentina, and parts of Bolivia––whereas *Calyptridium* and *Thingia* are North American, predominantly occurring in California and Oregon; *Cistanthe* is predominantly found in South America, but two species are North American (**Fig. 1**).

**Figure 1.**
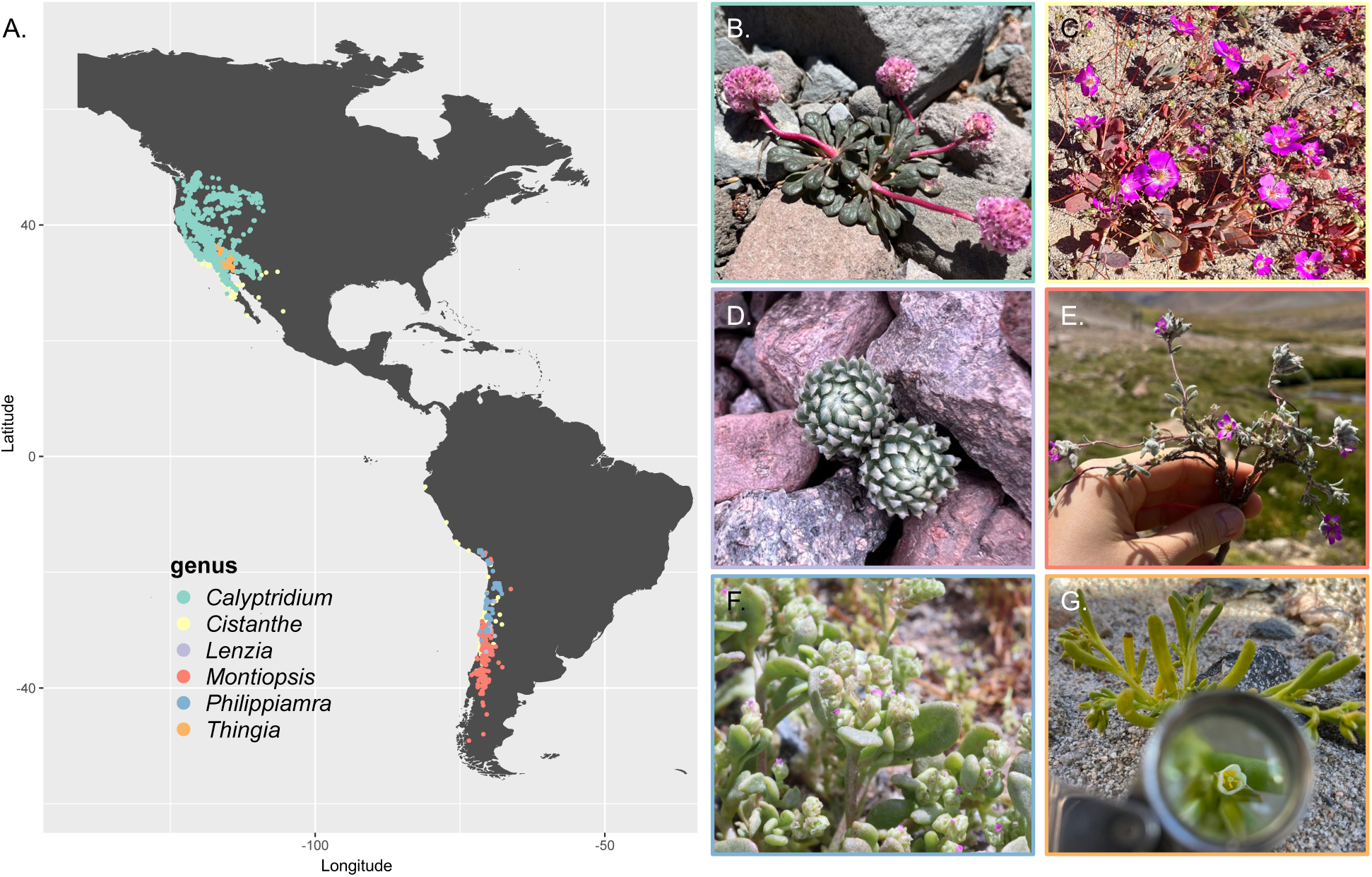
Geographic distribution of Cistantheae and a representative photo from each clade showing a diversity of growth forms and floral morphologies. **A.** Occurrence points on the map are filtered and cleaned data from the Global Biodiversity Information Facility (2022), and color coded by clade. From the top left: **B.** *Calyptridium umbellatum*, **C.** *Cistanthe aff. longiscapa*, **D.** *Lenzia chamaepitys*, **E.** *Montiopsis gilliesii*, **F.** *Philippiamra calycina*, **G.** *Thingia ambigua*.

The species in the Cistantheae are characterized by having dehiscent valvate capsules as fruit, ranging from one or few to numerous seeds per capsule. Although these herbaceous species exhibit a basic rosette body type (Ogburn and Edwards 2015), they vary considerably in habit; some are short-lived ephemerals, others are classical herbaceous perennials, while a few occasionally become shrubby and woody at the base. Many (though not all) of the species have succulent leaves (Ogburn and Edwards 2013); we know, for example, *Cistanthe grandiflora* can plastically increase their succulence (Ogburn and Edwards 2010). Additionally, multiple succulent species in the Cistantheae can facultatively utilize Crassulacean acid metabolism (CAM) photosynthesis upon drought (Arroyo *et al*. 1990; Holtum *et al*. 2021; Gilman *et al*. 2023; Chomentowska *et al*. 2025). Moreover, these species also exhibit substantial diversity in floral morphology, while all investigated species are self-fertile (Hinton 1976; Hinton 1976; Ford 1992). Ecologically, Cistantheae species are quite diverse, and occupy xeric, mesic/Mediterranean, and high-elevation montane habitats. The evolutionary transitions between these biomes appear to be correlated with transitions between annual (semelparous) and perennial (iteroparous) life history strategies (Ogburn and Edwards 2015). For example, *Philippiamra* species are desert-occupying annuals, surviving as seeds through prolonged drought in the Atacama Desert (Hershkovitz 2019), while some perennial *Montiopsis* species inhabit the Andean mountains up to 4000 m in altitude (Ford 1992; Peralta and Molina 2021).

Despite their ecological and morphological diversity, much uncertainty remains in Cistantheae taxonomy. Many of the species now classified in Cistantheae have been systematically moved out of *Calandrinia sensu lato* over time (Carolin 1987, 1993; Ford 1992; Cullen and Peralta 1994; Peralta 1988, 1993, 1999; Peralta and Ford-Werntz 2008; Hershkovitz 1990a,b, 1991, 1993a,b, 2018, 2019; Peralta and Molina 2021) after the floristic treatments of the 1800s (including Hooker and Arnott 1833; Barnéoud 1847; Philippi 1860, 1867, 1893; Howell 1893; Reiche 1897, 1898). Despite these efforts, decades of confusion persist and have been accompanied with many rounds of synonymizations and splitting of species (Franz 1908; Pax and Hoffmann 1934; Añón 1953; Kelley 1973; McNeill 1974; Hinton 1975; Nyananyo 1986, 1990; Carolin 1987, 1993; Ford 1992; Cullen and Peralta 1994; Peralta 1988, 1993, 1999; Ford-Werntz and Peralta 2002; Peralta and Ford-Werntz 2008; Guilliams 2009; Simpson *et al*. 2010; Guilliams *et al*. 2011; Guilliams and Miller 2014; Hershkovitz 1990a,b, 1991, 1993a,b, 2018, 2019; Watson 2019; JM Watson 2020; J Watson 2020; Peralta and Molina 2021). The difficulties may in part be due to the presence of intraspecific variation in morphology (e.g., in *Montiopsis*; Ford 1992), possibly due to rampant phenotypic plasticity. This is especially true in the desert taxa, where species are notably hard to identify (Hershkovitz 2019). Another source contributing to ambiguous species boundaries may be due to phenology, where many of the desert species in the Cistantheae will bloom synchronously post-rainfall event—a phenomenon commonly associated with El Niño years (Vidiella *et al*. 1999; Jaksic 2001; Chávez *et al*. 2019). Since many of these species live in sympatry (personal observation), the simultaneous flowering could facilitate easy transfer of pollen between different species and promote interspecific gene flow (Levin and Kerster 1974), obscuring traits associated with taxonomy or introducing hybrids (Hershkovitz 2006; Mark Hershkovitz 2019).

Importantly, the Cistantheae phylogeny, and especially the phylogenetic relationships within the subclade *Cistanthe*, remains essentially unresolved. No previous study has inferred a densely sampled phylogeny (i.e., including multiple specimens per species) for the group. Existing phylogenetic studies including Cistantheae have been either broader (suborder Portulacineae or the Montiaceae-wide) or focused on other Montiaceae clades (Hershkovitz and Zimmer 2000; Applequist and Wallace 2001; Hershkovitz 2006; Applequist *et al*. 2006; Guilliams 2009; Nyffeler and Eggli 2010; Price 2012; Ogburn and Edwards 2015; Goolsby *et al*. 2018; Hancock *et al*. 2018; Moore *et al*. 2018; Gilman *et al*. 2024), and often based on incomplete sampling, limited markers and accumulated GenBank sequences. In all, a robust phylogeny with adequate species representation is essential to clarify relationships in the clade, reexamine the taxonomy, and study patterns of trait evolution across Cistantheae.

In this study, we reconstructed the evolutionary history of the desert-alpine lineage Cistantheae using reduced-representation genome sequencing, including both reliably identifiable taxa as well as morphologically ambiguous specimens. We prioritized sampling species from the field (though our sampling was supplemented by herbarium specimens) and dense sampling within taxa to examine the monophyly of described species. We hoped to improve phylogenetic resolution in the backbone of Cistantheae and at shallow nodes, and to identify potential sources of incongruence (e.g., evidence of introgression, incomplete lineage sorting) that may partly explain taxonomic difficulties that persist. From previous phylogenetic work, we expected to find *Calyptridium*, *Philippiamra*, *Thingia* and *Lenzia* to be more closely related to each other than they are to *Montiopsis* and *Cistanthe* (Hershkovitz 2006; Guilliams 2009; Obgurn and Edwards 2015). We also anticipated the annual *Cistanthe* sect. *Rosulate* taxa from the Atacama Desert, a notoriously difficult taxonomic group, to be particularly recalcitrant to phylogenetic resolution.

With our new phylogeny, we re-evaluated the evolutionary correlations between life history strategy and climate niche. We hypothesized that annuality would be associated with warmer, drier, and lower-elevation habitats, whereas perennials would be associated with colder, less-dry, and higher elevation environments. Such a result would be consistent with what we find in the Montiaceae more broadly (where life history was correlated with temperature niche; Ogburn and Edwards 2015). However, this study includes more species of Cistantheae than this previous analysis. Therefore, Cistantheae (the clade with the highest lability in life history and climate niche transitions within Montiaceae; Ogburn and Edwards 2015) may alternatively deviate from the broader patterns with the inclusion of more taxa. Together, these analyses seek to clarify evolutionary relationships in a charismatic desert-alpine clade, and provide a well-supported phylogeny as the foundation for current and future taxonomic and comparative research in the Cistantheae.

## MATERIALS AND METHODS

### Data collection and ddRAD sequencing

We conducted multiple rounds of fieldwork between 2017-2023 in Peru, Chile, Argentina, and the USA to collect Cistantheae species (and other Montiaceae species to be used as outgroups). These voucher specimens were deposited at an herbarium of the specimens’ origin countries, including UNMSM Herbarium (SMF), Museo Nacional de Historia Natural (SGO), Herbario Ruiz Leal (MERL), Herbario Mendoza (MEN), Clifton Smith Herbarium (SBBG), as well as the Yale University Herbarium (YU). We also utilized dried leaf tissue or frozen DNA extractions from previously collected specimens (Arakaki et al. 2011; Ogburn and Edwards 2015), as well as herbarium specimens from Steere Herbarium (NY), San Diego State University Herbarium (SDSU), and Sukkulenten-Sammlung Zürich (ZSS). We extracted DNA from dried leaf tissue by first grinding them with TissueLyser III (Qiagen) with 5 mm stainless steel beads. We used a CTAB/β-mercaptoethanol-based extraction protocol and cleaned the extractions with Mag-Bind® Total Pure NGS (Omega Bio-tek) on magnetic racks. In all, we collected and extracted DNA from 164 samples (see **Supporting Information File 1** for the full list of specimens sampled).

Double-digest restriction site-associated DNA sequencing (ddRAD-Seq; Peterson *et al*. 2012) has been successfully applied in studies spanning population genomic and phylogenomic frameworks (e.g., Jacobs *et al*. 2021; MacGuigan *et al*. 2023; Anghel *et al*. 2025). We prepared ddRAD libraries from our extracted genomic DNA using a combination of PstI (rare cutter) and MspI (common cutter) restriction enzymes, as this enzyme-pair yields a high proportion of SNPs from genic and non-repeat regions of the genome in tomatoes (Shirasawa *et al*. 2016). Custom barcoded adapters were ligated to the digested DNA fragments such that samples could be multiplexed prior to PCR amplification. The amplified fragments were size-selected to retain those between 300-500 bp. The prepared ddRAD libraries were sequenced at the University of Oregon Genomics and Cell Characterization Core Facility (GC3F) on the Illumina HiSeq 4000 or NovaSeq 6000 machines (100-bp single-end reads).

### ddRAD sequence assemblies

The raw reads from the sequenced ddRAD libraries were demultiplexed and assembled using IPYRAD v.0.9.81 (Eaton and Overcast 2020). On average, ddRAD sequencing yielded *c.* 3,777,063 raw reads per sample. To minimize the risk of erroneously clustering paralogous loci (a worry with *de novo* assemblies), we opted for a reference-based assembly of the ddRAD sequences. Specifically, we implemented two separate assembly methods: one using the full *Cistanthe cachinalensis* genome (Chomentowska *et al*. 2025) as the reference, and another using only the coding-sequences from the same *C. cachinalensis* genome. Assembly parameters from IPYRAD included strict adapter filtering (filter_adapters = 2) and a maximum of 4 alleles per site to account for potential polyploidy (max_alleles_consens = 4). Otherwise, we applied slightly more conservative parameter values to reduce the instances of misaligning paralogs and to accommodate potentially messy alignments (e.g., limiting the proportion of shared polymorphic sites per locus with max_shared_Hs_locus = 0.25). Full parameter files and associated IPYRAD scripts are available at: https://github.com/anriiam/Cistantheae-phylogeny.

For the Cistantheae-wide phylogenetic analyses, we generated a final matrix based on the coding-sequence-referenced assembly (hereafter, CDS-referenced assembly) by subsampling 1–5 individuals per species, resulting in a total of 108 samples including outgroups. The whole-genome-referenced assembly (hereafter, genome-referenced assembly) contained too much missing data for further analyses at this broader scale. To balance loci retention with minimizing missing data, we required that each locus must be present in at least 30 samples in the final output (min_samples_locus = 30). We also produced assemblies for each major subclade—*Philippiamra*, *Calyptridium*, *Montiopsis*, *Cistanthe* sect. *Cistanthe*, and the combined *Cistanthe* sect. *Rosulatae* and sect. *Andinae—*including all sequenced individuals and varying the amount of missing data using the min_samples_locus parameter. Across subclade datasets, the genome-reference assembly yielded an average of ∼15378 consensus reads (putative loci) per sample (pre-filtering with across samples with the min_samples_locus parameter), while the CDS-referenced assembly yielded an average of ∼6514 consensus reads per sample. Full assembly statistics are available in **Supporting Information File 2.**

### Phylogenetic analyses

The final Cistantheae-wide CDS-referenced assembly contained 1,412 loci, with 80.18% missing data across the full sequence matrix of 263,221 sites (with 73.56% missing in just the SNP matrix containing 19,694 sites, of which 8,871 were parsimony-informative). Using this assembled matrix, we inferred a phylogeny with a maximum-likelihood approach implemented in IQ-TREE2 v.2.2.2 (Minh *et al*. 2020). We modified default parameters (-pers 0.2, -nstop 500) to better accommodate the characteristics of our dataset, which contains many short ddRAD sequences. To ensure that our IQ-TREE2 inferences were comparable across analyses, we used the GTR+I+G model of sequence evolution. We also verified that the resulting topology remained consistent when switching to MODELFINDER (the default model selection option), and when varying the starting tree—comparing results obtained using a random starting tree (-t RANDOM) versus a neighbor-joining tree (default settings). Support values were calculated using 1,000 ultrafast bootstrap (UFBoot) replicates.

At the subclade level, both the CDS-referenced assemblies and the genome-referenced assemblies were used to infer phylogenies. For phylogenetic inference based on the CDS-referenced alignments, we required that each locus be present in 30% of the samples within each subclade dataset; these assemblies contained between 43.61% and 58.21% of missing data. For the genome-referenced assemblies, we required that each locus be present in 90% of the samples for each subclade, resulting in 34.47% to 48.03% missing data. We prioritized including more data when feasible (which corresponds to more missingness but does not necessarily mean less informative; Eaton *et al*. 2017), but the genome-referenced assemblies produced the most sensical phylogenies when missing data was substantially reduced. We inferred maximum-likelihood trees using IQ-TREE2 as described above, and we also inferred coalescent-based quartet trees using the ‘tetrad’ module implemented in the IPYRAD-ANALYSIS toolkit (https://github.com/eaton-lab/tetrad) with 1,000 bootstrap replicates.

### Genetic structure and introgression

For analyses of genetic structure across all individuals in each subclade, we first filtered the VCF files from the subclade-level IPYRAD assemblies (min_samples_locus = 30%) using *VCFTOOLS* v.0.1.16 (Danecek *et al*. 2011). This included filtering out non-biallelic sites, as well as sites with minor allele frequencies less than 0.05 (--maf 0.05). Additionally, we thinned out the matrix by retaining one site every 700 base pairs to limit linkage among SNPs. We also filtered out genotypes with high missingness (--max-missing) requiring that 70–95% of individuals have a genotype called at a given SNP, while still retaining more than 1,000 SNPs per assembly.

These filtered assemblies were loaded into R v.4.4.0 (R Core Team 2024), where we assessed genetic structure and potential admixture by estimating ancestry coefficients using sparse nonnegative matrix factorization (sNMF), implemented in the package LEA v.3.16.0 (Frichot and François 2015); sNMF does not assume an evolutionary model, hence this was chosen above other popular methods like Structure. To determine the optimal number of genetic clusters (K), we used a cross-entropy criterion after testing K values from 2 to 20 (or 10, for smaller datasets) for each subclade. We used a regularization parameter α = 100 for smaller datasets, and ran each analyses with 100 replicates.

Furthermore, we wanted to detect instances of introgression using the ‘baba’ module in the IPYRAD-ANALYSIS toolkit, which calculates D-statistics (ABBA-BABA tests) from the .loci file outputs from IPYRAD (min_samples_locus = 30%). For each analysis, we used either the subclade-level maximum likelihood or Tetrad trees to draw all possible four-taxon trees to generate tests. A D-statistic value greater than zero indicates an excess of the ‘ABBA’ allelic pattern, while a value less than zero indicates an excess of a ‘BABA’ allelic pattern (where A = ancestral and B = derived; Durand *et al*. 2011). We particularly focused these analyses on samples that showed mixed ancestry in the sNMF results, changed position across different tree inferences, or were suspected of hybrid origin due to morphology. Statistical significance was determined using Z-scores calculated for each test, with |Z| > 3 considered significant.

### Trait evolution and correlation

In order to assign life history strategy to each species (annual or perennial), we used a combination of expert knowledge and descriptions from protologues and floras. To obtain geographical ranges and construct species climate envelopes, we downloaded and filtered occurrence data from the Global Biodiversity Information Facility (https://GBIF.Org 2025); these occurrence points were used to extract 19 bioclimatic variables and elevation from WorldClim v.2 (Fick and Hijmans 2017), which were then averaged for each species. The number of GBIF occurrence records per species ranged from 2 (*Philippiamra pachyphylla*) to 1,354 (*Calyptridium monandrum*), with an average of 91.2 specimens per species. To estimate the elevational range for each species, we calculated interquartile range (IQR), which ignores outliers; we used linear models to test whether mean elevation predicts elevational range within annual and perennial species separately, as well as whether the relationship differs between the two life history strategies.

We trimmed the Cistantheae-wide phylogeny to include just one sample per species. For instances where a species was nested within another, both taxa were kept for this analysis (effectively reflecting a sister relationship between those species). We did, however, opt to collapse the *Cistanthe longiscapa* complex, where there was no phylogenetic resolution among samples (see Results), to one tip (*Cistanthe longiscapa;* this includes individuals we had morphologically assigned to *C. litoralis* and *C. cymosa*). We time-calibrated the trimmed tree using secondary calibrations from Arakaki *et al* (2011). This was done using the R package PHYTOOLS v.2.6.5 (Revell 2024), specifically the chronos() function which uses penalized likelihood. We calibrated three nodes (the Montiaceae crown, the node where *Calandrinia* + *Lewisiopsis* and Cistantheae diverge, and the node where *Cistanthe paniculata* and the rest of *Cistanthe* sect. *Cistanthe* diverge), and estimated divergence times for all other nodes using 10 discrete rate categories under a molecular clock (model = “discrete”) with a smoothing parameter (lambda = 0.1).

We conducted a series of phylogenetic comparative analyses to test the correlation between life history strategy and climate variation across Cistantheae. To explore the difference in climatic niche between annuals and perennials, we first conducted a principal components analysis (PCA) using the PCA() function in the FACTOMINER v.2.11 package (Lê *et al*. 2008). To infer the ancestral states of life history strategies, we used the ancr() function implemented in PHYTOOLS (Revell 2024). We fit both the equal-rates (ER) and the all-rates-different (ARD; effectively an asymmetric model with a binary trait) models via the fitMk() function, and estimated ancestral states as weighted averages (using an ANOVA-based model comparison) between the two models. For continuous climate variables and elevation, we used the fastAnc() function in PHYTOOLS to reconstruct ancestral states under a Brownian motion model. Specifically for elevation, we visualized raw GBIF occurrence points using ridge plots with the R package GGRIDGES (Wilke 2024), retaining only the portion of each density curve above a minimum relative height of 0.01.

Finally, we calculated trait correlations between life history and each continuous climate variable using multiple approaches. First, we performed phylogenetic generalized least squares (PGLS) regression using pgls() from the CAPER v.1.0.3 R package (Orme *et al*. 2023), treating life history as a binary predictor (0 = annual, 1 = perennial) and each climate or elevation variable as the response. To account for the independent comparisons of multiple traits, we applied a false discovery rate (FDR) correction (Benjamini and Hochberg 1995) to the *p*-values. We also used PGLS to test whether elevational range differs significantly between annual and perennial species. Second, we used the phylolm() function from the PHYLOLM package (Ho and Ane 2014) to fit an Ornstein–Uhlenbeck (OU) model of trait evolution (model = “OUfixedRoot”). For each trait, we compared the fits of the OU and Brownian motion models using Akaike information criterion (AIC), and extracted the regression coefficients from the preferred model. Third, we used the threshold model implemented in the threshBayes() function in PHYTOOLS (Revell 2024) to jointly model the evolution of life history liability and continuous climate variables under Brownian motion. The liability of a binary trait (in our case, life history strategy) can be interpreted as the underlying/unobserved continuous values that determine whether a species is more likely to be an annual or perennial, given a threshold value (Felsenstein 2005, 2012). We ran MCMC chains for 1 million generations, discarding the first 10% as burn-in. Convergence was assessed by examining the posterior density plots and calculating the effective sample size (ESS). We used the posterior distributions to estimate the evolutionary correlation (posterior sample mean) between life history and a given climate variable (deemed significant if the 95% credible interval excluded zero).

## RESULTS

### Phylogenetic inference of Cistantheae

The representative ddRAD phylogeny of the Cistantheae contained 108 specimens, which included 7 outgroup species as well as 48 Cistantheae species, which represent ∼63% of the described Cistantheae taxa. Of these, 10 are *Calyptridium*, five are *Philippiamra*, 14 are *Montiopsis*, and 17 are *Cistanthe*, as well as the monotypic *Lenzia chamaepitys* and *Thingia ambigua*. The resulting Cistantheae maximum-likelihood phylogeny shows well-supported relationships among the major subclades (**Fig. 2**).

**Figure 2.**
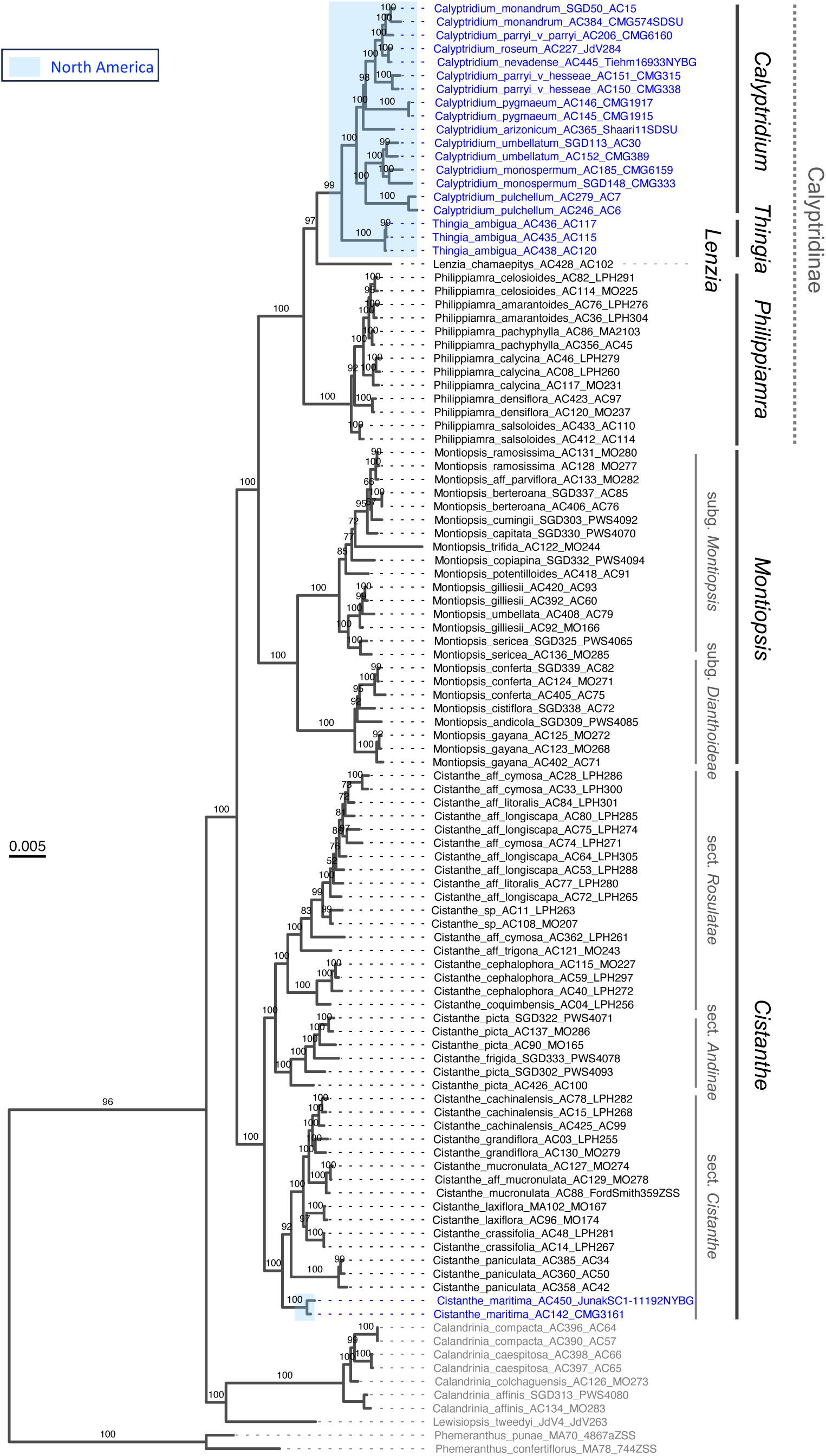
Maximum-likelihood phylogeny of Cistantheae using coding-sequence referenced ddRAD assembly. The subclades are annotated on the right. The tree was inferred using IQ-TREE2 with 1,000 Ultrafast Bootstraps (support values on branch), and includes 108 samples. Outgroup taxa are indicated by gray tip labels. In the ingroup, the North American clades are emphasized with blue tip labels, as well as with a light blue highlight around the clades on the phylogeny. Otherwise, all the rest are Peruvian, Chilean, and/or Argentinean taxa.

The topology places *Cistanthe* as sister to the rest of the clade—contrary to previous inferences which placed *Cistanthe* and *Montiopsis* as sister lineages (Ogburn and Edwards 2015). Instead, *Montiopsis* appears more closely related to the Calyptridinae Hershk. (which includes *Philippiamra*, *Lenzia*, *Thingia*, *Calyptridium*) than to *Cistanthe* (also inferred in Hancock *et al*. 2018). The monotypic *Lenzia* is sister to the *Thingia* + *Calyptridium* clade, and *Philippiamra* the rest of Calyptridinae (**Fig. 2**). The only nodes along the backbone with less than 100% UFBoot support are the split between *Lenzia* and the *Thingia* + *Calyptridium* clade (UFBoot = 97), and the split between *Thingia* and *Calyptridium* (UFBoot = 98). Because of the limitation to interpreting ultrafast bootstrap values on concatenated phylogenomic datasets, these two splits should be considered only weakly supported.

This tree also supports at least two long-distance, amphitropical dispersal events in Cistantheae (**Fig. 2**). The first occurs along the branch leading to the *Thingia* + *Calyptridium* clade, which is composed of North American species and is sister to the perennial, high-elevation Andean *Lenzia*—which, as stated before, are all sister to *Philippiamra*, a clade made entirely up of arid-adapted annuals. The second instance involves *Cistanthe maritima*, a North American desert species that is sister to the rest of *Cistanthe* sect. *Cistanthe*, which are all South American.

### Subclade-level phylogenetic analyses

Subclade-level analyses provided greater resolution and support for certain species relationships across datasets and inference methods, as compared to the broader Cistantheae-wide tree. While some taxa were problematic, there was no consistent pattern that could explain differences in topology between datasets that exclude and include non-coding regions, or between concatenation and species-tree approaches.

#### Calyptridium

This clade is well sampled, and includes 10 of 11 described taxa (minimum-rank). The relationships among the North American pussypaws (*Calyptridium*) are also mostly resolved. Across both the Cistantheae-wide and subclade-level phylogenies, the *Calyptridium umbellatum* complex (including perennial *C. umbellatum*, perennial *C. monospermum*, and annual *C. pulchellum*; *sensu* Hinton 1975; Guilliams 2009) forms a clearly distinct, independently evolving lineage, while the remaining *Calyptridium* species (all annual) form a separate clade (**Figs 2, 3A, 4A; Sup. Figs 1, 2**).

**Figure 3.**
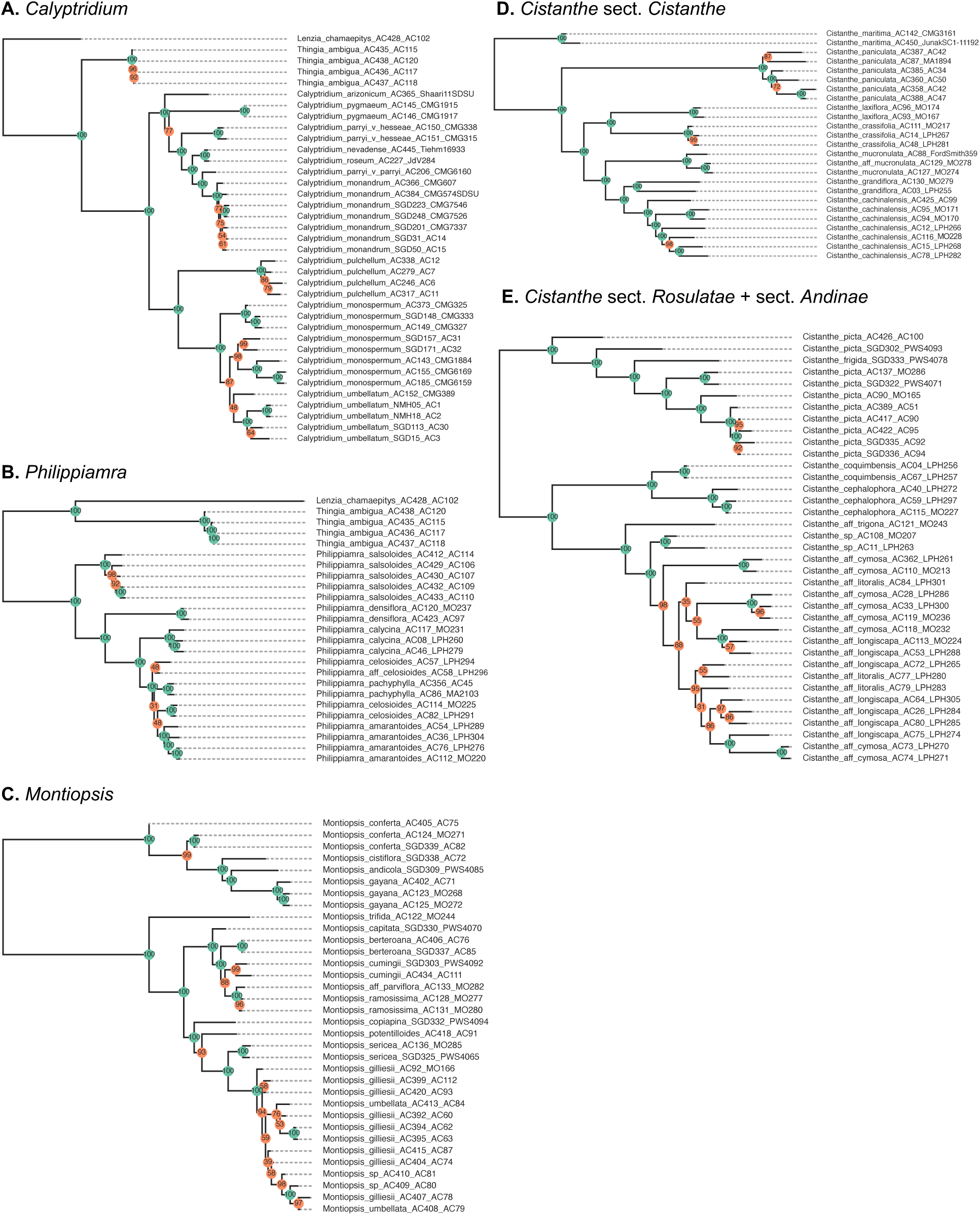
Maximum-likelihood (ML) phylogenies using whole-genome-referenced ddRAD assemblies in **A.** *Calyptridium*, **B.** *Philippiamra*, **C.** *Montiopsis*, **D.** *Cistanthe* sect. *Cistanthe*, and **E.** *Cistanthe* sect. *Rosulatae* and sect. *Andinae*. The trees were inferred using IQ-TREE2 with 1,000 Ultrafast Bootstraps, with support values on nodes (support = 100 in green; support < 100 in orange). Subclade-level ML phylogenies using coding sequence-referenced ddRAD loci are available in the Supporting Information.

We should note, however, that neither *C. umbellatum* nor *C. monospermum* are monophyletic in the more densely sampled subclade-level phylogenies. Furthermore, the relationships among the samples of these two species differ between the whole-genome-referenced and coding-sequence-referenced trees (**Figs 3A, 4A**). There is strong evidence of introgression between three of the *C. monospermum* samples (CMG325, CMG327, CMG333) and other *Calyptridium* species: specifically, CMG327/CMG333 with the other annual *Calyptridium* samples (**Fig. 5A**), and CMG325 with *C. pulchellum* (**Fig. 5B**). CMG325 is particularly noteworthy, since the sample represents a herbarium specimen collected from a *C. monospermum* population found outside of its typical range. In the coding-sequence-referenced maximum likelihood tree and both Tetrad trees, one *C. umbellatum* sample (CMG389) falls outside of the otherwise monophyletic *C. umbellatum* samples. CMG389 also shows evidence of introgression with the annual *Calyptridium* species (**Fig. 5C**).

**Figure 4.**
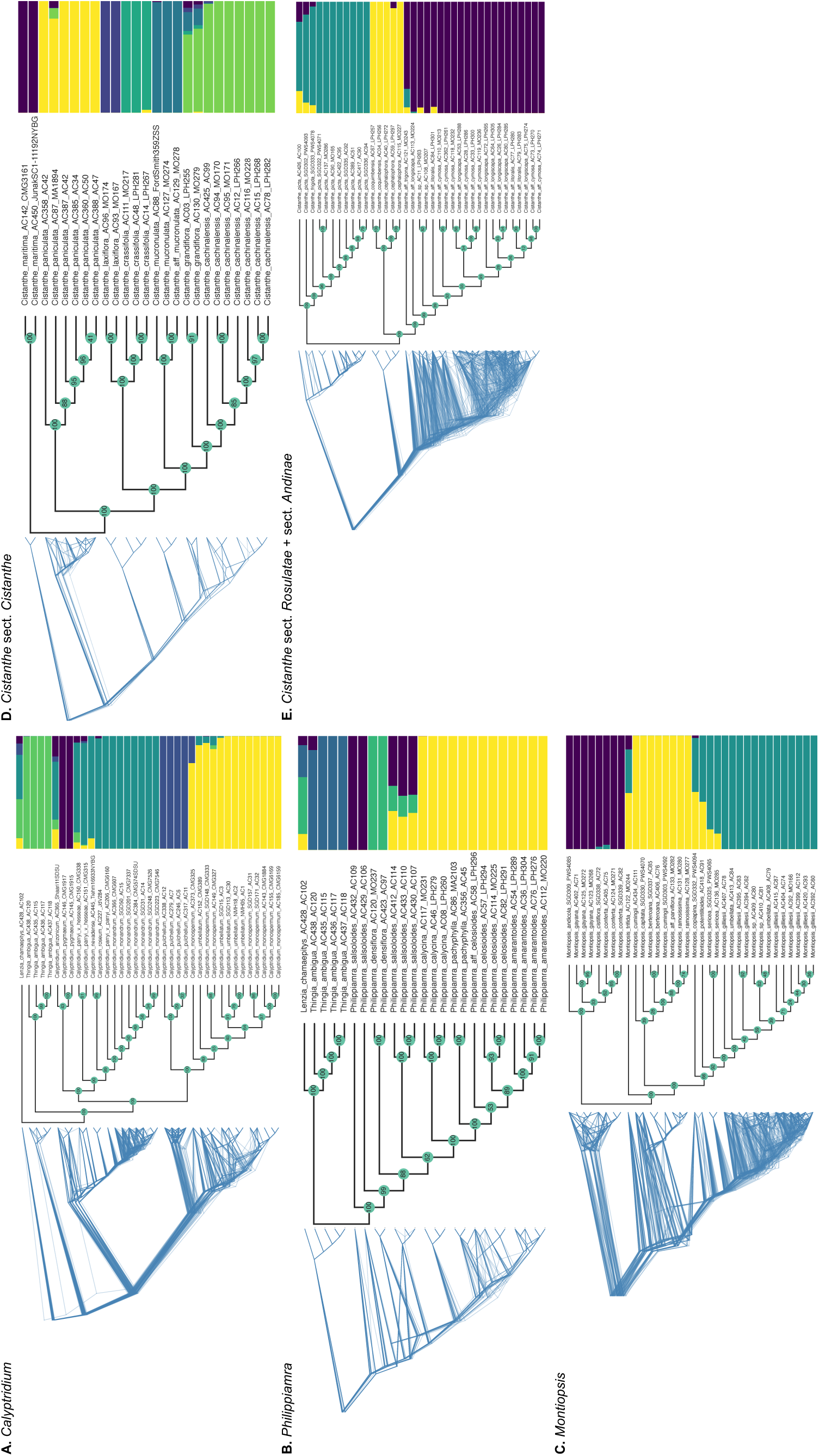
Tetrad phylogenies (coalescent-based quartet trees) and genomic structure using whole-genome-referenced ddRAD assemblies in **A.** *Calyptridium*, **B.** *Philippiamra*, **C.** *Montiopsis*, **D.** *Cistanthe* sect. *Cistanthe*, and **E.** *Cistanthe* sect. *Rosulatae* and sect. *Andinae*. For each subclade panel, the leftmost tree shows a sample of 100 out of 1,000 quartet-based bootstrap replicates, and the middle tree shows the consensus Tetrad phylogeny. The rightmost graph shows genetic clustering for each subclade, as inferred by ancestry coefficients estimated with sparse nonnegative matrix factorization (sNMF). The colors represent the optimal number of genetic clusters (K) for each subclade (**Sup. Fig. 7**). Each horizontal bar represents an individual sample (in the same order as the tips on the phylogenies), and the proportions within each bar correspond to the estimated ancestry coefficients. Same analyses using coding-sequence-referenced ddRAD assemblies for each subclade are available in the Supporting Information.

**Figure 5.**
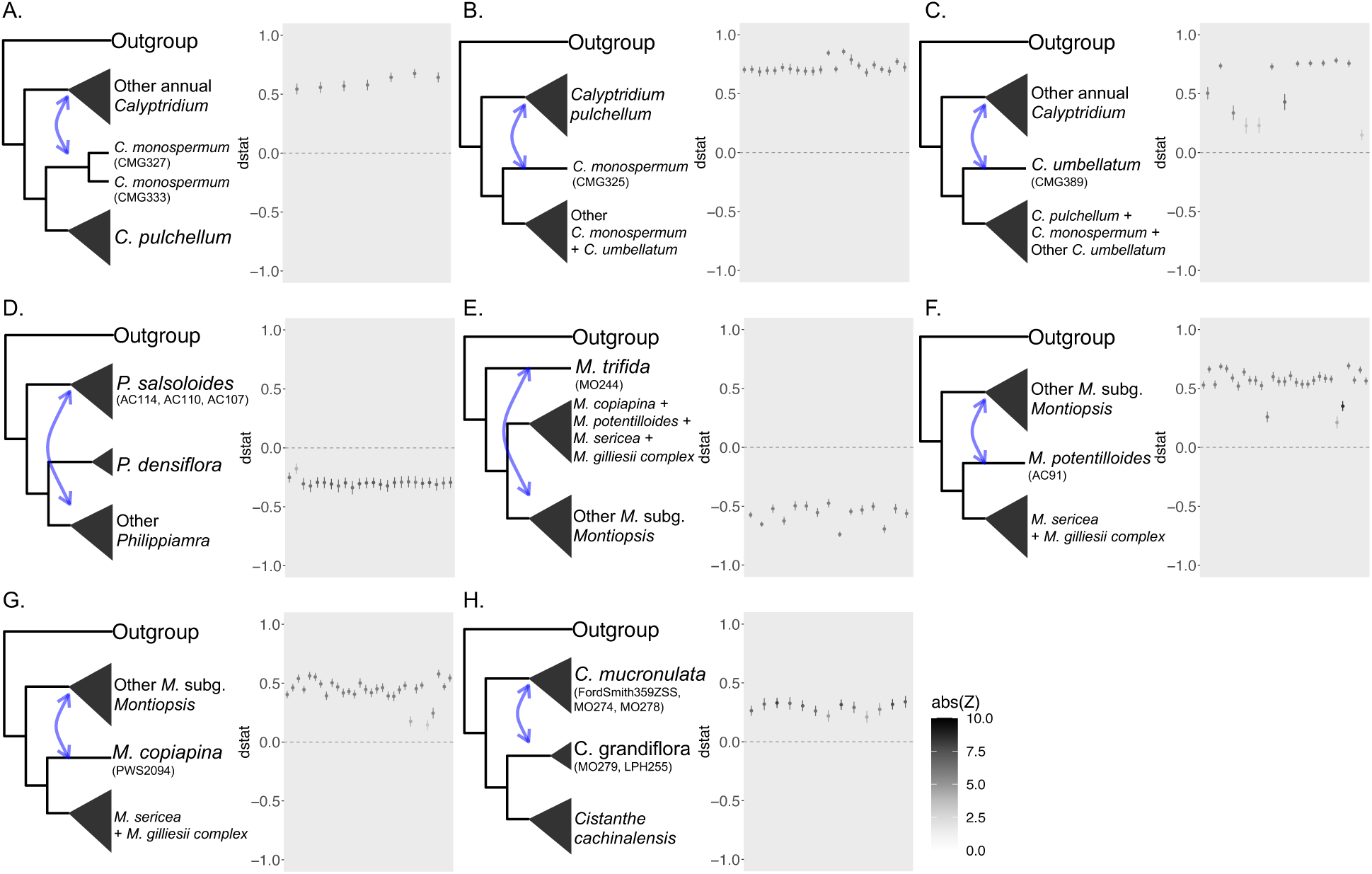
Results from ABBA-BABA tests using the whole-genome-reference loci, indicating introgression. In each panel, the cladograms on the left illustrate the test setup; in Newick format, the test follows the topology (Outgroup, (P3, (P2, P1)), where tips may represent individual samples or clades. Each point in the graphs on the right side of the panels represent the mean *D*-statistic from 100 repetitions per test, with 95% confidence intervals shown. The tests consist of every possible permutation of the P2 and P1 taxa. Positive *D-*statistics indicate excess allele sharing between the P3 and P2 taxa, whereas negative values indicate excess sharing between P3 and P1 (as summarized by the blue arrows on the cladograms). While all *D*-statistic values shown here are statistically significant (|Z| >3), the color intensity of each point corresponds to the magnitude of |Z|. Panels A., B. and C. are ABBA-BABA tests in *Calyptridium*; D. *Philippiamra*; E., F. and G. *Montiopsis*; panel H. *Cistanthe*.

Within the annual *Calyptridium* clade, all minimum-rank taxa are monophyletic, except for *C. parryi*; however, it is worth noting that several of these annual species are currently represented by only a single sample (**Figs 3A, 4A**). The non-monophyly of the previously described *C. parryi* varieties is consistent with results from a previous five-gene phylogeny (Guilliams 2009). These consistent results support the relatively new combination of *C. hesseae* (J.T. Thomas) Hershk. and *C. nevadense* (J.T. Thomas) Hershk. as their own separate species from the basionyms where they were described as varieties of *Calyptridium parryi* (Hershkovitz 2019), though this conclusion would be strengthened with the inclusion of additional samples for each variety. Most importantly, our trees support the conclusions of Simpson *et al*. (2010) which established *C. arizonicum* as its own species based on morphological differences from *C. parryi* and other previous varieties. Relationships among these species in the clade are inconsistent from earlier works; specifically, in both Guilliams (2009) and Ogburn and Edwards (2015), *C. pygmaeum* and *C. roseum* are sister taxa. Nevertheless, the phylogenies presented here boast mostly strong support for each split (**Figs 2, 3A, 4A; Sup. Figs 1, 2**).

#### Philippiamra

This clade is well sampled, and includes five of seven described species. The phylogenetic relationships among the extremely arid-adapted annuals in *Philippiamra* are relatively straightforward, with little to no incongruence between the full Cistantheae tree (**Fig. 2**) and the subclade-level trees (**Figs 3B,4B; Sup. Figs 1, 2)**. The species relationships are generally well-supported and morphologically recognizable. Notably, the Chilean *Philippiamra celosioides* is more closely related to the Chilean *P. amarantoides* than it is to the Peruvian *P. pachyphylla*, despite past taxonomic treatments that synonymized *P. pachyphylla* with *P. celosioides* (Govaerts *et al*. 2021).

In the subclade-level analysis, we had two additional samples of *P. celosioides* (LPH294 and LPH296). LPH294, a morphologically distinct and sterile sample, formed a monophyletic group with the other two *P. celosioides* samples in the genome-referenced Tetrad tree (**Fig. 4B**), as well as the coding sequence-referenced ML (**Sup. Fig. 1**) and Tetrad trees (**Sup. Fig. 2**); LPH296 on the other hand, was outside of this three-sample clade. In the genome-referenced ML tree (**Sup. Fig. 3B**), however, LPH294 and LPH296 were sister to each other and not monophyletic with the other *P. celosioides* samples. The behavior of LPH296 could be due to introgressive gene flow or admixture, which we found a strong signal for with *P. calycina* (**Fig. 5D**). All of the *P. celosioides* samples do belong in a strongly supported monophyletic group with *P. pachyphylla* and *P. amarantoides* samples—a pattern that was recovered across datasets and inference methods (**Figs 3B, 4B; Sup. Figs 1, S2**). However, the support values for the relationship between these species were low across trees. In all, the monophyly of *P. pachyphylla* and *P. celosioides* remain questionable, especially given that *P. amarantoides* appears nested within this group with strong support. Finally, there is some discordance in the two Tetrad trees (**Fig. 4; Sup. Fig. 2**) where *P. salsoloides* samples are not monophyletic, likely due to shared derived alleles (via introgression) between three specific *P. salsoloides* samples (AC412, AC433, and AC430) and the *P. calycina*/*P. pachyphylla*/*P. celosioides*/*P. amarantoides* clade (**Fig. 5D**).

#### Montiopsis

This clade is well sampled, and includes 14 of 19 described species. *Montiopsis* species, many of which occur at higher elevations than most other Cistantheae species, clearly fall into two clades considered as two subgenera: *Montiopsis* subg. *Dianthoideae* and *Montiopsis* subg. *Montiopsis* (Ford 1992). The former consists of only perennial species, while the latter includes both annuals and perennials. Across the different phylogenies, support for species relationships within each of the two clades ranges from strong to weak (**Figs 2, 3C, 4C; Sup. Figs 1, 2**). However, sampling remains limited in some of these species, several of which are represented by only one sample.

In the full Cistantheae tree, *Montiopsis gayana* is sister to the rest of the species in subg. *Dianthoideae*, though the relationships among the three remaining species are weakly supported (**Fig. 2**). In contrast, the subclade-level maximum likelihood trees show a well-supported topology: *M. gayana* is instead sister to *M. andicola*, which are together sister to *M. cistiflora,* which are all nested within a non-monophyletic *M. conferta* (**Fig. 3C; Sup. Fig. 1**). Further, there was no evidence of introgression among these samples, suggesting that the observed incongruences likely stem from incomplete lineage sorting across the subclade (Goulet *et al*. 2017). Morphologically, species in subg. *Dianthoideae* have tricolpate pollen, whereas the species in subg. *Montiopsis* have pantoporate pollen, complex barbellate trichomes, and three-toothed sepals (Ford 1992; Peralta and Molina 2021).

On the other hand, there is much less resolution among species in subg. *Montiopsis*, particularly in the Cistantheae-wide tree (**Fig. 2**). Still, the sister-relationship between the perennial *M. sericea* and the perennial *M. gilliesii* complex is well-supported. We found that *M. umbellata* samples were non-monophyletic and instead nested among *M. gilliesii* samples leading us to treat this as a species complex. This complex includes *Montiopsis sp.* AC410 and AC409 which we suspected were hybrids of *M. gilliesii* and *M. umbellata* in the field based on their morphology. Removing these ‘hybrid’ samples did not change the non-monophyly of *M. gilliesii* and *M. umbellata*. The two other perennial species, *M. potentilloides* and *M. copiapina*, subtend the annuals in subg. *Montiopsis* with minimal support. There is low support for species relationships among the annuals in the Cistantheae-wide tree (**Fig. 2**). Moreover, *M. potentilloides*, *M. copiapina*, and *M. trifida* exhibit mixed ancestry across genetic clusters (**Fig. 4C**) and may have experienced recent introgressive gene flow (**Figs 5E,F,G**).

The relationships within subg. *Montiopsis* are better resolved in the coding-sequence-referenced subclade tree (**Sup. Fig. 3**), which is largely consistent with the Cistantheae-wide tree topology with the exception of the strongly supported sister-relationship between *M. trifida* and *M. capitata*. However, the topology shifts considerably in the genome-sequence referenced subclade tree, where *M. trifida* is inferred as sister to the rest of subg. *Montiopsis* species, and *M. potentilloides* and *M. copiapina* are sister to the other perennials (*M. sericeae* and the *M. gilliesii* complex; **Fig. 3C**). Together, the evidence of mixed ancestry and topological discordance point to a complex evolutionary history that has likely shaped the diversification of subg. *Montiopsis*.

#### Cistanthe

Cistanthe is the most species-rich clade in Cistantheae, and also the group in most need of taxonomic work. We sampled only 17 of 37 described species. In the Cistantheae-wide phylogeny, *Cistanthe* species form three subclades; one with species that have mostly been assigned to *Cistanthe* sect. *Cistanthe* Reiche (with the exception of *C. maritima*), one with perennials in *Cistanthe* sect. *Andinae* Reiche, and one with desert annuals included in *Cistanthe* sect. *Rosulatae* Hershk (Hershkovitz 2019). We inferred *C. maritima*, a North American species, as sister to the rest of *Cistanthe* sect. *Cistanthe* (composed entirely of South American species) with strong support. Within the South American clade, *C. paniculata* (a Peruvian species) is sister to the rest—a relationship that has weak support in the Cistantheae-wide tree (**Fig. 2**) but strong support in the subclade-level trees (**Figs 3D, 4D; Sup. Figs 1, 2**). This is consistent with previous inferences from Smith *et al*. (2018) and Hancock *et al*. (2018). Moreover, the inferred sister-species relationship between *C. crassifolia* and *C. laxiflora* has strong support at the subclade-level trees (**Figs 3D, 4D; Sup. Figs 1, 2**). The relationships among the remaining *Cistanthe* sect. *Cistanthe* species are strongly supported in both the broader (**Fig. 2**) and subclade level phylogenies (**Figs 3D, 4D; Sup. Figs 1, 2**). All the species in this subclade are perennial except for two: *Cistanthe maritima*, subtending the rest of sect. *Cistanthe*, and *C. cachinalensis*, nested well within the group. Additionally, we found evidence of introgressive gene flow between *C. mucronulata* and *C. grandiflora* (**Fig. 5H**).

In contrast, the phylogenetic inference in the rest of *Cistanthe* is far less straightforward. *Cistanthe sect. Andinae*, which is represented by the high-elevation perennial species *C. picta* and *C. frigida* in this phylogeny, is sister to the rest of the desert annuals, with 100% UFBoot support in the Cistantheae-wide tree (**Fig. 2**). Furthermore, *C. frigida*—a species found at some of the highest elevations within the Cistantheae—is nested within *C. picta*. Notably, our *C. picta* sampling straddles the Chile-Argentina border, on either slope of the western Cordillera of the Andes. In all the subclade-level trees (**Figs 3E, 4E; Sup. Figs 1, 2**)., the Chilean *C. picta* samples form a gradient (including *C. frigida*) before the monophyly of the Argentinian *C. picta* samples (MO165, AC51, AC90, AC95, AC92, AC94), suggesting some geographical structure to the topology.

After the divergence of the *C. picta* clade, a well-supported clade of *C. coquimbensis* and *C. cephalophora* is inferred as sister to the rest of *Cistanthe* sect. *Rosulate* (across all the phylogenies; (**Figs 2, 3E, 4E; Sup. Figs 1, 2**). This clade corresponds to the taxonomic group ‘subsect. *Thyrsoideae*’ from Hershkovitz (2019). The rest of sect. *Rosulate* is poorly resolved, and we will refer to it here as the ‘*C. longiscapa* complex’. All the specimens in this clade are united by conspicuous shiny petals, though within the clade, no morphologically delineated species appear to be monophyletic. *Cistanthe longiscapa* is the iconic species known for painting the Atacama Desert magenta pink during ‘superblooms’.

In the *C. longiscapa* complex, *C.* aff. *trigona* (MO243) is consistently inferred as sister to the rest of the samples in the group, which forms a comb-like topology in the Cistantheae-wide tree (**Fig. 2**). Still, the subclade-level tree from coding sequence-referenced loci shows stronger support and some emerging patterns (**Sup. Figs 1, 2**). Two *Cistanthe* sp. samples (MO207 and LPH263) resemble the vegetative morphology of *C. arenaria* (but are clearly annuals with smooth seeds), and diverge from the rest of the complex with strong support. Though they were collected many years apart and by different collectors, they are geographically close, and might represent two individuals of the same population. Within the remainder of the complex, we then see a split between two subgroups. Both subgroups include samples with *C. cymosa*-like inflorescence morphology and samples with *C. longiscapa*-like traits (**Sup. Fig. 3**). According to previous descriptions, *C.* aff. *cymosa* samples (as defined by the inflorescence morphology) should have smooth seeds, whereas *C.* aff. *longiscapa* samples should have pusticulate-tomentose seed morphology (Kelley 1973). However, we found both morphotypes with both seed types. There are *C.* aff. *cymosa* and *C.* aff. *longiscapa* samples with smooth seeds (the latter, therefore, may more appropriately be referred to as *C.* aff. *litoralis*; Hershkovitz 2022), and *C.* aff. *cymosa* and *C.* aff. *longiscapa* samples with pusticulate-tomentose seeds (**Fig. 3**). One sample, LPH305, even exhibited hairy seeds reminiscent of *C. cachinalensis*. The overall topology of the complex is broadly similar in the genome-referenced tree at the subclade-level, though the support values are starkly lower (**Figs 3E, 4E**). An interactive map of the specimens in the sect. *Rosulatae* tree alongside the genome-referenced Tetrad phylogeny demonstrates specimen clustering and geography: https://camayal.info/wa/treetom/?id=5Z1rykPR29ChQAG8QzSj (via TREETOM, Maya-Lastra 2020). The distribution of the samples does not suggest that different morphologies in the complex could be explained by geography; for example, *C.* aff. *litoralis* localities span Parque Nacional Llanos de Challe to the city of Taltal, with various *C.* aff. *longiscapa* and *C.* aff. *cymosa* specimens found in between. Geography does explain many of the phylogenetic groupings and sister-relationships between specimens, but cannot *completely* explain them either; for example, *C.* aff. *cymosa* MO214 and LPH261 are from different sites.

### Evolutionary correlation between life history and environmental variables

We found that life history is evolutionarily labile across the Cistantheae (**Fig. 6A**). Ancestral state reconstruction using the ancr() function from PHYTOOLS suggests that the ancestral life history state is ambiguous, though a perennial ancestor has a higher marginal probability. However, this result may be biased by the fact that all the sampled outgroup taxa in the tree happen to be perennial. Using a threshold model to estimate the liability across ancestral nodes yielded a similar result: the estimated liability of the Cistantheae ancestor was close to zero (i.e., near the threshold). This suggests uncertainty in the ancestral life history state, though with a slight preference for perenniality based on the posterior distribution. Assuming a perennial ancestor, annuality appears to have evolved independently at least five times within the clade. We also infer two subsequent shifts from annual to perennial—once along the *Lenzia* branch, and again in the lineage leading to *Calyptridium umbellatum* and *C. monospermum*.

**Figure 6.**
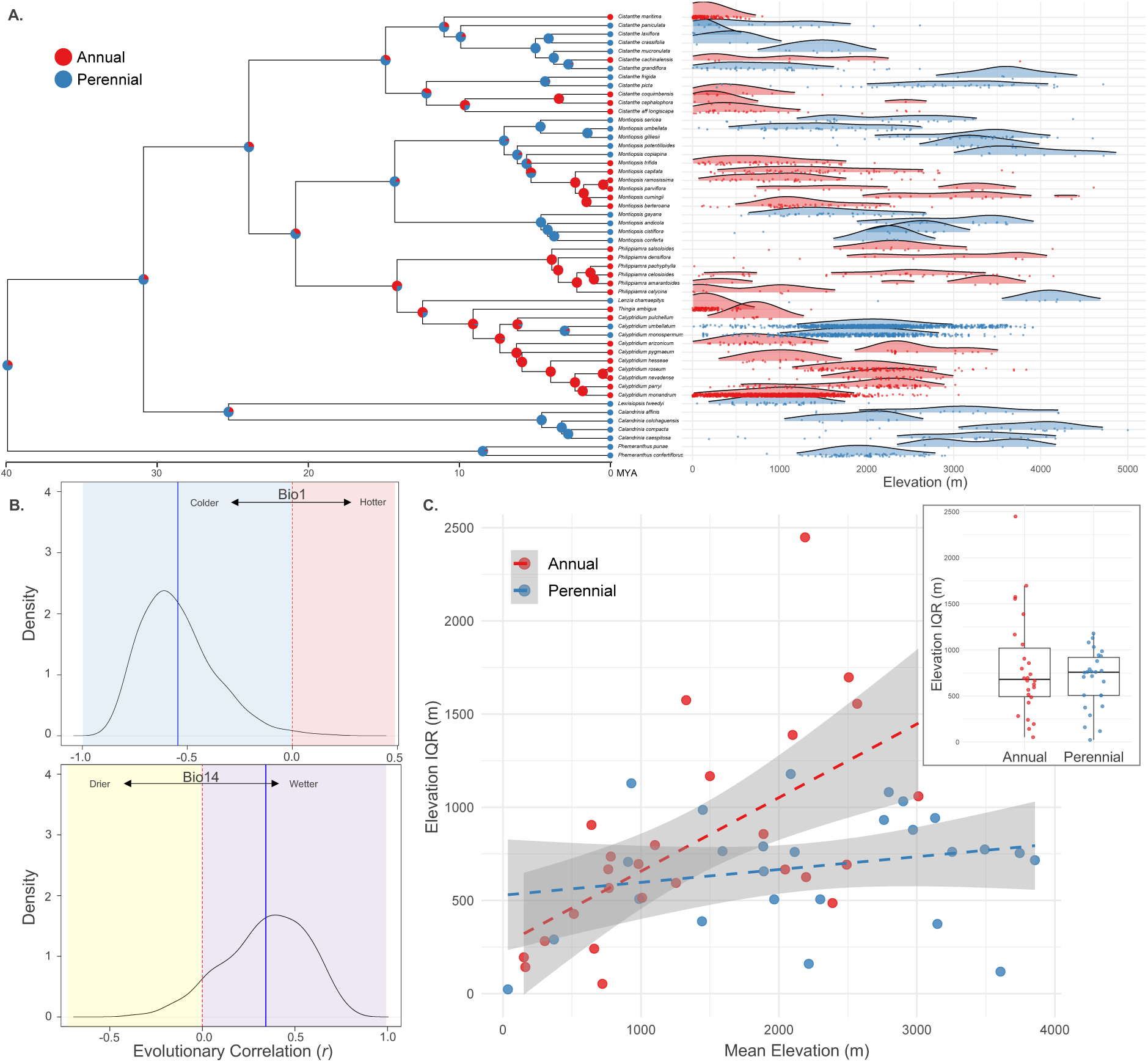
Life history and climate niche evolution across Cistantheae. **A.** Ancestral state reconstructions of life history strategy and corresponding species-level elevation distributions. The phylogeny shown (left) is time-calibrated and trimmed to one sample per species (see **Sup. Fig. 8**). Tip colors indicate life history strategy: red for annual species and blue for perennials. Pie charts at internal nodes represent the inferred likelihood of each ancestral state (annual versus perennial), following the same color scheme. Elevation ridge plots (right) use the same color scheme, with each point representing an occurrence record from cleaned Global Biodiversity Information Facility (2022) data. Density curves were visualized with the minimum relative height of 0.01. **B.** Posterior density distributions of evolutionary correlations (*r*) between life history and two climate variables: Bio1 (Mean Annual Temperature, top panel) and Bio14 (Precipitation of Driest Month, bottom panel). Vertical red dashed lines denote *r* = 0; solid blue lines denote the posterior mean. Positive correlations indicate that perennials tend to occur in colder (Bio1) and wetter (Bio14) environments. **C.** Linear regression of mean elevation versus elevation interquartile range (IQR) by life history strategy. Each point represents a species; red and blue points represent annuals and perennials, respectively. A significantly positive relationship in annuals indicates that annual species with higher mean elevations tend to have broader elevational ranges (*p* = 0.0011). This slope differs significantly from the perennial slope (*p* = 0.0077). The boxplot inset (top right) comparing IQR between annuals and perennials shows no statistically significant differences in elevational range between life history strategies.

We also found bioclimatic variables to be quite labile across the phylogeny (**Sup. Fig. 4**). A PCA of climate values (averaged per species) showed moderate separation between annuals and perennials, though with some overlap (**Sup. Fig. 5**). Patterns of trait correlations between these climate variables and life history were largely consistent with our expectations: across the phylogeny, perennials are associated with colder, wetter, and higher-elevation environments, whereas annuals tended to occupy hotter, drier, and lower-elevation habitats (**Table 1**; **Figs 6A,B; Sup. Fig. 5**). Furthermore, there was no significant difference in elevation interquartile range (IQR) between life history strategies (**Fig. 6C**). However, a linear regression of mean elevation versus elevation IQR showed a significant positive relationship in annuals (*p* = 0.0011), with no corresponding trend in perennials (**Fig. 6C**): these two slopes were significantly different (*p* = 0.0077). Moreover, the number of specimens per species is not a significant predictor of elevation IQR (*p* = 0.879), indicating sampling effort does not bias our estimates of elevational range. Across all traits, OU models consistently fit the data better than BM models, based on ΔAIC values from phylolm (**Table 1**).

**Table 1.**
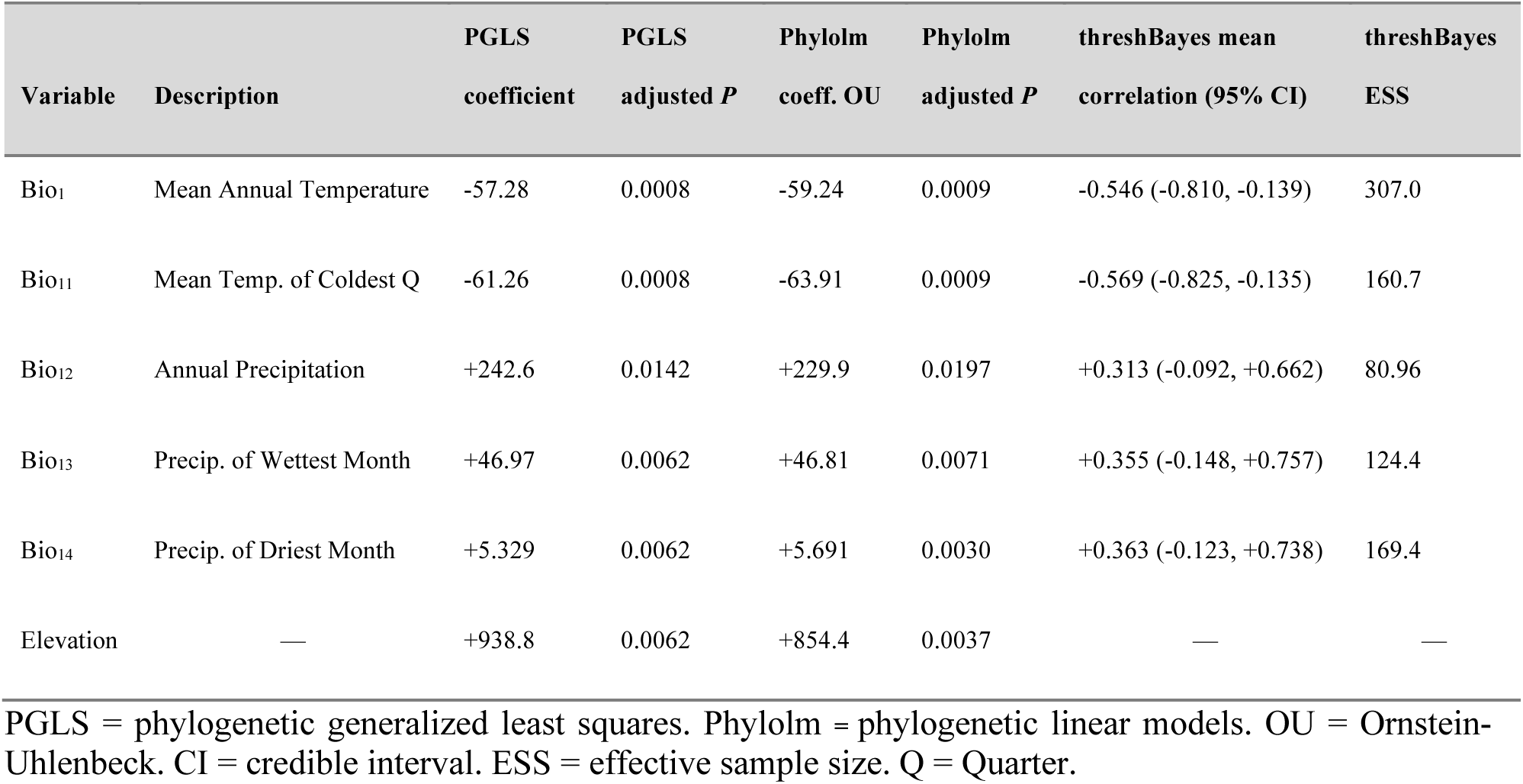
Phylogenetic correlations between environmental variables and life history in the Cistantheae.

Specific temperature-related variables were significantly and strongly correlated with life history strategy across the phylogeny. Based on PGLS (using the pgls function in CAPER) and phylogenetic linear models assuming an OU process (PHYLOLM, model = “OUfixedRoot”), the strongest signals were found among Bio11 (mean temperature of coldest quarter), Bio1 (mean annual temperature), Bio6 (minimum temperature of coldest month), and Bio10 (mean temperature of warmest quarter). Large negative regression coefficients were consistently recovered across both analyses (**Table 1; Sup. Table 1**). For example, Bio11 had a coefficient of –61.26 (*p* = 0.0008) based on the PGLS analysis, and a coefficient of –63.91 (*p* = 0.0002) in the OU phylogenetic linear model (**Table 1**). Such regression-based results were also supported by Bayesian threshold model analyses using threshBayes from PHYTOOLS. The estimated posterior mean correlations were also consistent with the direction of regression coefficients from PGLS and phylolm analyses (**Table 1**). For example, Bio1 had a posterior mean evolutionary correlation coefficient of -0.546 (95% credible interval between -0.810 and -0.139) with an effective sample size (ESS) of 307.8, which confirms convergence (**Fig. 6B**). Taken together, these analyses suggest that perennials were more associated with lower temperatures across the tree.

Precipitation-related variables were also correlated with life history strategy across the tree, though the associations were generally weaker. For example, Bio12 (annual precipitation), Bio13 (precipitation of wettest month), and Bio14 (precipitation of driest month) were positively correlated with perenniality in the PGLS analyses (Bio12 coefficient = +242.6, *p* = 0.00922; Bio13 coefficient = +46.97, *p* = 0.00277; Bio14 coefficient = +5.329, *p* = 0.00261; **Table 1**), suggesting that perennials tend to occur in wetter environments. These results were consistent in both direction and magnitude with results from OU phylogenetic linear models (**Table 1**). The threshBayes analyses showed similar directional trends in the correlation coefficient, supporting the association of wetter habitats with perenniality. However, for these precipitation-related variables like Bio 14 (ESS = 420.9; **Fig. 6B**), the 95% credible intervals included zero, which means the correlation coefficients were not statistically significant even if the directionalities were consistent (**Sup. Fig. 6**). Elevation also showed a moderate positive correlation with life history across the tree (PGLS coefficient = +911, p = 0.0018; phylolm coefficient = +876, p = 0.0015), suggesting that perennials tend to occupy at higher elevations than annuals.

## DISCUSSION

The Cistantheae phylogeny presented here (**Fig. 2**) represents a significant step forward in our efforts to develop the Montiaceae as a ‘model clade’ (*sensu* Donoghue and Edwards 2019)—a monophyletic group with a manageable number of species, allowing researchers to rigorously study any and all traits in a phylogenetic framework to ultimately infer underlying mechanisms of evolution. We are focusing on one tractable subclade at a time (e.g., Cistantheae in this current study; *Parakeelya* in Hancock *et al*. 2018) working toward a completely sampled species-level phylogeny of Montiaceae.

In many ways, the Cistantheae represent much of the ecological and morphological diversity that is found in the Montiaceae more broadly. Ecologically, many of these species have an affinity to montane and high-elevation habitats or mediterranean climates, while other xeric species will flower synchronously in the Atacama ‘superblooms’ to paint the desert magenta-pink (Chávez *et al*. 2019). Cistantheae is yet another example of a lineage traversing desert and alpine habitats—like *Linanthus* (Bell and Patterson 2000; Anghel *et al*. 2025), *Cirsium* (Kelch and Baldwin 2003; Siniscalchi et al. 2023), and *Oxalis* (Heibl and Renner 2012)—but exhibits an especially labile ecological niche, even within Montiaceae (Ogburn and Edwards, 2015). Physiologically, both perennial and annual *Cistanthe* species in the clade can facultatively upregulate CAM photosynthesis, an adaptation to drought that confers increased water use efficiency, when subjected to drought conditions (Holtum *et al*. 2021; Chomentowska *et al*. 2025). Cistantheae species also present a gradient of leaf succulence (Ogburn and Edwards 2013), a trait that is related to the expression of CAM (Gilman *et al*. 2024). Biogeographically, we found at least two independent long-distance dispersal events in the Cistantheae, which are both examples of an American amphitropical disjunct (AAD) dispersal pattern; this adds to the 237 ADD divergence events that fit this amphitropic pattern of distribution found so far in vascular plants (Simpson *et al*. 2017).

### Phylogenetic analyses of Cistantheae

Overall, our efforts have clarified the backbone of Cistantheae, providing strong support for the major clades and relationships between them. There has been substantial taxonomic work in this group over the last 50 years, including Carolin (1987; 1993), Kelley (1973), Hershkovitz (1990a,b, 1991, 1993a,b, 2018, 2019), Peralta (1993; 1999), Ford (1992), Ford-Werntz and Peralta (2002), Peralta and Ford-Werntz (2008), Guilliams (2009), Simpson *et al*. (2010), Guilliams and Miller (2014), and Peralta and Molina (2021). These studies are good progress from the treatments of the late 1800s (Hooker and Arnott 1833; Barnéoud 1847; Philippi 1860, 1867, 1893; Howell 1893; Reiche 1897, 1898), but they were largely undertaken without a strongly supported molecular phylogeny.

The overall topology of the Cistantheae phylogeny highlights *Montiopsis* as a particularly interesting clade. In Ogburn and Edwards (2015), *Montiopsis* was inferred as sister to *Cistanthe* (based on five markers). Hancock *et al*. (2018), on the other hand, inferred different relationships depending on marker sampling: *Montiopsis* was sister to *Cistanthe* in the 115-loci tree, whereas *Montiopsis* was sister to Calyptridinae in the 297-loci tree. Our phylogeny infers the latter relationship (**Fig. 2**). These topological differences may reflect conflicting evolutionary histories among different genes or ancient introgression at the origin of *Montiopsis*. Furthermore, our phylogeny places *Lenzia* sister to *Thingia* + *Calyptridium*, which are all then most closely related to *Philippiamra*, which matched our expectations (**Fig. 2**). However, this topology should be confirmed in future work, specifically because the support is low in the node that unites *Lenzia* with the *Thingia* + *Calyptridium* clade.

Within each of these clades, the relationship between recognized taxonomic groups were also clarified. By and large, the taxonomic assignments of species were recapitulated in the phylogeny (making up subclades), and the relationships between these subclades were also generally well-supported. For example, *Montiopsis* consists of subclades that correspond to subgenera *Montiopsis* and subgenera *Dianthoideae*; *Cistanthe* includes sections *Cistanthe*, *Rosulatae*, and *Andinae* with the latter two subclades as sister (**Fig. 2**). However, within each of these subclades there is persistent uncertainty about many species relationships and species delineation, highlighting the need for further investigation.

### *Cistanthe longiscapa* complex and evolution in the desert

Although a taxonomic revision of the *Cistanthe longiscapa* complex is clearly needed, defining species boundaries may prove difficult in these types of complexes with potentially rampant gene flow, incomplete lineage sorting (**Fig. 4E**), and phenotypic plasticity. Our results also suggest that the vegetative, inflorescence, and seed morphology characters often used to distinguish species in this complex may not be reliable taxonomic indicators.

Moreover, the complexity of phenology in the Atacama Desert must be taken into account when considering the evolution of this clade. The desert species—particularly those in the *Cistanthe longiscapa* complex— are often associated with ‘superblooms’ during El Niño years, which are characterized by above-average precipitation. While this association is generally true, these species can also flower in drier years, though germination and growth is quite patchy across the landscape. Even in years with substantial rainfall, blooms can remain patchy and unpredictable; the same site may be completely barren in one El Niño year despite having been in full bloom during another, while a field just nearby may show the opposite pattern. What are the consequences of such patchiness and episodic reproduction on the rate of evolution and patterns of gene flow? While no single model currently captures all the interacting forces that shape diversification in desert plants, there are relevant empirical studies that speak to individual components. For example, Falahati-Anbaran *et al*. (2014) found that gene flow through time (i.e., many generations of seeds germinating simultaneously) can counteract genetic variability introduced to a population via dispersal from another population in *Arabidopsis thaliana*.

It appears that in the *Cistanthe longiscapa* complex, putative species can easily reproduce with each other, and interspecific gene flow is rampant, which further muddies morphological differentiation (and taxonomic resolution). With the addition of the temporal and geographical separation of flowering and stochastic gene flow between populations, the complex has not sorted into independently evolving lineages that we can easily define as species. What might be more helpful in understanding how evolution works in this group is a population-level view into how SNPs, geography, phenology, and morphology interact. Assigning species to organize or categorize diversity between populations is *useful* (Levin 1979), but it also may not be appropriate in certain population biology settings, especially in plants (Raven 1976). While we aim to eventually sample all described species in *Cistanthe*, we predict that many will fall into the *C. longiscapa* complex and warrant a careful taxonomic revision.

### Future work in the *Calyptridium umbellatum* complex

Another area for future work lies in the underexplored diversity of the *Calyptridium umbellatum* complex, particularly in the perennial species. In the subclade-level analyses with all the voucher specimens that were available, the perennial *C. umbellatum* is nested within *C. monospermum* (the other perennial in *Calyptridium*). While the two species overlap in range, *C. umbellatum* also occurs at higher latitudes. We may find genetic or morphological differentiation that might correspond to appropriate species boundaries within the current circumscriptions of *C. monospermum* or the wider *C. umbellatum* complex—more so if we collect from populations at higher latitude where we were unable to sample for this study. It may be worthwhile to revisit the varietal designations in the *C. umbellatum* complex from Hinton (1975) based on morphology. Additionally, applying multispecies coalescent models (Jackson *et al*. 2017; Leaché *et al*. 2019) may be helpful for species delimitation in this complex.

### Implications of current gaps in taxon sampling

As previously discussed, our phylogeny includes only a single sample of *Lenzia*, and its placement remains uncertain. Although we recovered the relationship where *Lenzia* is sister to the *Thingia* + *Calyptridium* clade, support for this node is low, and its long branch may be driving spurious relationships via long-branch attraction. A denser sampling of *Lenzia* (though they are quite rare plants with a small geographical range in high-elevation Andes, making it a hard species to collect), and the use of whole-genome sequencing may help to clarify its phylogenetic position. It is also imperative to include *Hectorella* and *Lyallia*, two monospecific taxa currently not classified in the Cistantheae (but rather as Hectorelleae Appleq., Nepokr. & W.L. Wagner; Hershkovitz 2019) but with some morphological similarity to *Lenzia* with wildly disjunct distributions in the Kerguelen Islands and New Zealand, respectively. It is possible that *Lenzia* is actually sister to the rest of Calyptridinae (as inferred by previous studies by only a handful of markers and with weak support; Hershkovitz 2006; Ogburn and Edwards 2015).

In addition, several other key taxa remain unsampled in our current phylogeny. For example, the North American *Cistanthe guadalupensis* was excluded due to poor DNA extraction quality. This species is most likely sister to *C. maritima* (*C. guadalupensis* has been shown or hypothesized to be sister to species in *Cistanthe* sect. *Cistanthe*; Applequist and Wallace 2001; Hershkovitz 2019), though we cannot currently rule out the possibility of yet another independent amphitropical dispersal event. Otherwise, the species relationships within the rest of *Cistanthe* sect. *Cistanthe*—to which, we found, *C. maritima* is sister—are well-resolved and strongly supported. In *Cistanthe* sect. *Rosulate*, however, we are missing notable perennials such as *C. arenaria* and *C. fenzlii*, which are likely close relatives of the annual *C. longiscapa* complex (Hershkovitz 2022b). Including these taxa may reveal an additional transition between annual and perennial life history strategies. Lastly, there are several described species in the *C. longiscapa* complex that were not included in our analyses; future work will determine whether these additional morphological varieties are embedded within the complex or represent independently evolving lineages (*sensu* De Queiroz 2007).

### ddRADSeq: its advantages and limitations

Double-digest RAD sequencing (ddRADSeq) has remained popular over the past decade for good reason: it is a cost-effective method of generating many non-random genomic fragments across many individuals (Peterson *et al*. 2012). One of the strengths of ddRADSeq is its sequencing depth-to-cost ratio, allowing application at both phylogenomic and population-genomic scales (e.g., Psonis *et al*. 2021; Lavretsky *et al*. 2021; Anghel *et al*. 2025). However, our study also encountered the limitations of ddRADSeq for phylogenetic inference. As with many reduced-representation approaches, missing data was substantial and very few loci were supporting certain splits. Our findings suggest that we may be approaching the upper limit of ddRADSeq’s utility for deeper divergences in fast-evolving herbaceous lineages (as opposed to slower-evolving woody clades, but see Vargas *et al*. 2017). In contrast, ddRADSeq had more power (in the form of more loci and more complete matrices) in discerning species-level relationships in the subclade analyses. Even then, coalescent-based inferences showed considerable discordance among ddRAD loci, which in part explains why there is little phylogenetic resolution in certain areas like the *C. longiscapa* complex (**Fig. 4E**). We also detected evidence of introgression and admixture, which may be contributing to the taxonomic uncertainty and phylogenetic conflict in this clade. However, these analyses can be used with a variety of data types such as whole-genome skimming at shallower depth, which has become much more economical. Future studies of Cistantheae population structure and phylogenomics would likely benefit from transitioning to alternative sequencing methods for greater applicability and improved resolution for both shallow and deeper evolutionary inference.

### Life history evolution in Cistantheae

Beyond clarifying subclade and species relationships, this Cistantheae phylogeny in its current form is also useful in exploring patterns of trait evolution. Particularly, we examined how life history strategy might be related to climatic variables. Unlike previous studies, which relied on one sample to represent a given species and few molecular markers, our phylogeny is based on many newly collected and sequenced specimens per species and thousands of loci. Our results were broadly consistent with patterns identified across Montiaceae (Ogburn and Edwards 2015): perennial species tend to occupy colder climates while annuals tend to occupy warmer (**Table 1**). This suggests that evolutionary correlations between life history strategy and temperature niche persist across phylogenetic scales, at least in lineages where life history remains evolutionarily labile across the trees at different scales. We also found a weaker, but still notable, association between life history and precipitation, where perennials tend to be occupying wetter habitats. These results align with those from other plant lineages, highlighting the potential roles that temperature and aridity play in shaping life history evolution (Evans *et al*. 2007; Datson *et al*. 2008; Wang *et al*. 2016; Monroe *et al*. 2019; Boyko *et al*. 2023).

Why might life history strategies be strongly correlated with environmental variables, across diverse lineages and phylogenetic scales? For one, the life history strategies represent different ways plants may allocate resources and biomass under specific environmental constraints. In deserts where precipitation is sporadic and drought stress is imminent, high seedling survival is more advantageous compared to low adult mortality (Charnov and Schaffer 1973; Stearns 1992). So, based on these trade-offs, annuals should be favored in more unpredictable and harsher climates (Stearns 1976). However, similarly ‘harsh’ alpine or high-elevation habitats rarely host annual plants—likely due to shorter growing seasons and lower temperatures (relative to deserts) that limit a plant’s ability to complete their life cycle (Körner and Larcher 1988; Körner 2003). Under these conditions, low adult mortality and perenniality may instead be more advantageous. Moreover, different morphological and physiological traits are related to plant life history strategies as well (Roumet *et al*. 2006; Lundgren and Des Marais 2020). For example, high allocation to below ground biomass may be considered a part of a perennial ‘syndrome’, a trait especially important for surviving in higher elevations and its associated climate (Lundgren and Des Marais 2020). These traits associated with life history strategies evolve repeatedly and correlatively (comprising ‘life history syndromes’) with environmental niche: perennial species being more common in colder, wetter, and higher-elevation environments, while annuals being favored in warmer, drier, low-elevation habitats (Hjertaas *et al*. 2023).

However, the exceptions to these patterns are especially interesting. Both annual and perennial *Cistanthe* species exist in desert environments, often in sympatry; the same is true for the annual and perennial *Montiopsis* species in mid-elevational bands in the Andes. Interestingly, the architecture of desert perennials is more diverse than alpine/high elevation perennial *Cistanthe*, which are herbaceous perennials with extensive belowground woody root systems. This growth form also exists in the desert (*C. arenaria*, *C. fenzlii*, *C. grandiflora*), but there are also species with persistent aboveground tissue, becoming soft-wooded shrubs (*C. laxiflora* and relatives). In Eriogonoideae, Kostikova *et al*. (2013) found that perennials had broader niche breadth than annual species, even while occurring at higher mean elevations. Contrastingly, climatic niches tend to evolve more rapidly in herbaceous lineages than woody ones (Smith and Beaulieu 2009), and faster molecular evolution in annuals may further facilitate this adaptability (Yue *et al*. 2010). Cistantheae thus provides an excellent system to explore evolutionary shifts in life history, particularly in relation to species that occupy narrow niche space versus those that span broad climate and elevational gradients (i.e., an ‘ecological generalist’). Such ecological flexibility may be key to evolution of certain traits; for example, a study of pollination syndromes in *Roscoela* has found that generalized pollinators, rather than just specialized-pollinators, are important for the evolution of pollinator-related traits in flowers (Paudel *et al*. 2019). In Cistantheae, the prevalence of ecological generalists supports the idea of reciprocal pre-adaptation to both alpine and desert environments. Across the phylogeny, many Cistantheae species exhibit broad elevational niches that may be exemplary of adaptability in both directions (**Fig. 6A**). This is perhaps facilitated by climatic overlaps between these environments, such as similarly low relative humidity (Billings and Mooney 1968; Ehleringer 1985; Körner 2003). Although perennials tend to occur at higher elevations and annuals at lower, some sister species with the same life history strategy show little overlap in their elevation ranges (**Fig. 6A**). In general, elevational niche appears to be evolutionarily labile across the Cistantheae phylogeny. While there is no significant difference in the elevation interquartile range (IQR) between life history strategies, the species with the broadest elevational ranges are annuals— potentially reflecting greater ecological plasticity or dispersal potential in these species (**Fig. 6C**). Significantly, annuals exhibit a significant positive relationship between mean elevation and elevation IQR, which suggests that annuals with higher mean elevations tend to occupy broader elevational niches. In general, Cistantheae species demonstrate broad elevational tolerances, which would provide a diversity of ecological selection pressures that might in turn facilitate repeated evolutionary shifts between life history strategies.

Lastly, there is a continuum between annual and perennial life history strategies, which introduces further complexity in the discussion of life history evolution. Some annuals are facultatively perennial under favorable environmental conditions; conversely, some perennials can behave as annuals under disturbance or water limitation (Friedman 2020). In the Cistantheae, *Cistanthe cachinalensis* has been categorized as an annual, but has also been described as a biennial (Philippi 1860; Reiche 1989). In our experiences growing *C. cachinalensis* in the growth chamber, they can become somewhat woody at the base of the rosette when grown under well-watered conditions; interestingly, some genotypes have multiple bouts of reproduction, suggesting the genes underlying perenniality may be partially intact, whereas others clearly senesce after one round of flowering (personal observations). This species belongs to an otherwise perennial *Cistanthe* sect. *Cistanthe* lineage, representing a recent transition from perenniality to annuality (**Fig. 6A**). Therefore, it is perhaps unsurprising that *C. cachinalensis* could variably behave as a facultative perennial. In contrast, perenniality has recently (re-)evolved in *Calyptridium monospermum* and *C. umbellatum* in an otherwise annual *Calyptridium* lineage—another subclade suited for further investigation. We are starting to understand differences in the genetic mechanisms underlying investment into flowering versus vegetative growth between annuals and perennials (e.g., in *Mimulus*, Gould *et al*. 2017; or Brassicaceae, Albani and Coupland 2010; Kiefer *et al*. 2017). Furthermore, many such genes are likely pleiotropic (Friedman 2020). Future research could combine target analyses of candidate genes with comparative methods to explore the evolution of life history strategies in Cistantheae.

## Conclusions

This study presents the first densely sampled, high-throughput molecular phylogeny of Cistantheae. We clarify key phylogenetic relationships and identify long-distance dispersal events, abundant instances of admixture, and new species complexes that warrant further attention. Comparative analyses reveal a consistent association between life history strategies and environmental variables: annuals are associated with hotter, drier, lower-elevation environments, while perennials were associated with cooler, wetter climates. Yet, the most compelling insights come from the exceptions to these trends—including broad ecological ranges and facultative behaviors—which accompany the evolutionary lability of life history strategies. Rather than representing fixed sets of traits and characteristics, annuality and perenniality appear to encompass complex and dynamic syndromes in Cistantheae, and perhaps other plant lineages, which merit further investigations.

## ACKNOWLEDGEMENTS

We would like to thank and acknowledge Thomas Near, Carlos Maya-Lastra, Jose Moreno-Villena, Karolina Heyduk, Edgar Benavides, Morgan Moeglein, Ava Ghezelayagh, Liam Taylor, Alison Carranza for training, help, and advise on sequencing and analyses. We also thank Samantha O’Brien and Sarah Teichman with substantial help in the field. We would like to additionally thank Michael Donoghue, Susan Mazer, and Ian Gilman for valuable insights on the overall project. A special thank you to UNMSM Herbarium, Museo Nacional de Historia Natural, Herbario Ruiz Leal, Clifton Smith Herbarium, and Yale University Herbarium, Steere Herbarium, San Diego State University Herbarium, and Sukkulenten-Sammlung Zürich for their collaboration and assistance. Funding for this project was provided by the US National Science Foundation Division of Environmental Biology grant No. 2327957, and the Sussex Fund through the Yale Peabody Museum.

## Author contributions

AC, AMM, CMG, EJE, IEP, JAMH, LG, LPH, NH, PWS, RMO, and SGD conducted fieldwork and collected plant specimens and tissues. AC, NH and SGD extracted DNA, and AC prepared genomic libraries. AC assembled the ddRADSeq data, inferred phylogenies, and conducted downstream analyses. AMM, CMG, EJE, IEP, and PWS provided essential taxonomic expertise. AC and EJE prepared the manuscript, while all authors edited or reviewed and approved the manuscript.

## Data availability

All raw, demultiplexed sequencing data will be made available on NCBI Sequence Read Archive (SRA). Alignment files, tree files, character and climate matrices, and scripts used for analyses are available at https://github.com/anriiam/Cistantheae-phylogeny.

## SUPPORTING INFORMATION

**Sup. Fig. 1.** Subclade-level Maximum-Likelihood (ML) phylogenies using coding sequence-referenced ddRAD assemblies.

**Sup. Fig. 2.** Subclade-level Tetrad phylogenies (coalescent-based quartet trees) and genomic structure using coding sequence-referenced ddRAD assemblies.

**Sup. Fig. 3.** Distribution of morphological characters in the *Cistanthe longiscapa* complex.

**Sup. Fig. 4.** Ancestral state reconstructions of bioclimatic variables across Cistantheae.

**Sup. Fig. 5.** PCA of bioclimatic variables.

**Sup. Fig. 6.** Posterior density distributions of evolutionary correlations between life history and climate variables across Cistantheae.

**Sup. Fig. 7.** Cross-entropy estimates from sparse nonnegative matrix factorization (sNMF) analyses for each subclade.

**Sup. Fig. 8.** Time-calibrated phylogeny of representative Cistantheae species.

**Sup. Table 1.** Summary statistics from phylogenetic generalized least squares (PGLS) and Ornstein-Uhlenbeck phylogenetic linear models (phylolm) between life history and bioclimatic variables.

Following supporting files are available on https://github.com/anriiam/Cistantheae-phylogeny.

**Sup. File 1**: List of vouchers and specimens sampled for phylogenetic analyses.

**Sup. File 2**: Assembly statistics from ddRAD assemblies.

