## Supplementary material for "Phylogenomic analyses of the diverse desert-alpine plant lineage Cistantheae": Sup.

The following Supporting Information is available for this article:

**Sup. Fig. 1.** Subclade-level Maximum-Likelihood (ML) phylogenies using coding sequence-referenced ddRAD assemblies.
**Sup. Fig. 2.** Subclade-level Tetrad phylogenies (coalescent-based quartet trees) and genomic structure using coding sequence-referenced ddRAD assemblies.
**Sup. Fig. 3.** Distribution of morphological characters in the *Cistanthe longiscapa* complex.
**Sup. Fig. 4.** Ancestral state reconstructions of bioclimatic variables across Cistantheae.
**Sup. Fig. 5.** PCA of bioclimatic variables.
**Sup. Fig. 6.** Posterior density distributions of evolutionary correlations between life history and climate variables across Cistantheae.
**Sup. Fig. 7.** Cross-entropy estimates from sparse nonnegative matrix factorization (sNMF) analyses for each subclade.
**Sup. Fig. 8.** Time-calibrated phylogeny of representative Cistantheae species.

**Sup. Table 1.** Summary statistics from phylogenetic generalized least squares (PGLS) and Ornstein-Uhlenbeck phylogenetic linear models (phylolm) between life history and bioclimatic variables.

**Sup. File 1:** List of vouchers and specimens sampled for phylogenetic analyses.
**Sup. File 2:** Assembly statistics from ddRAD assemblies.

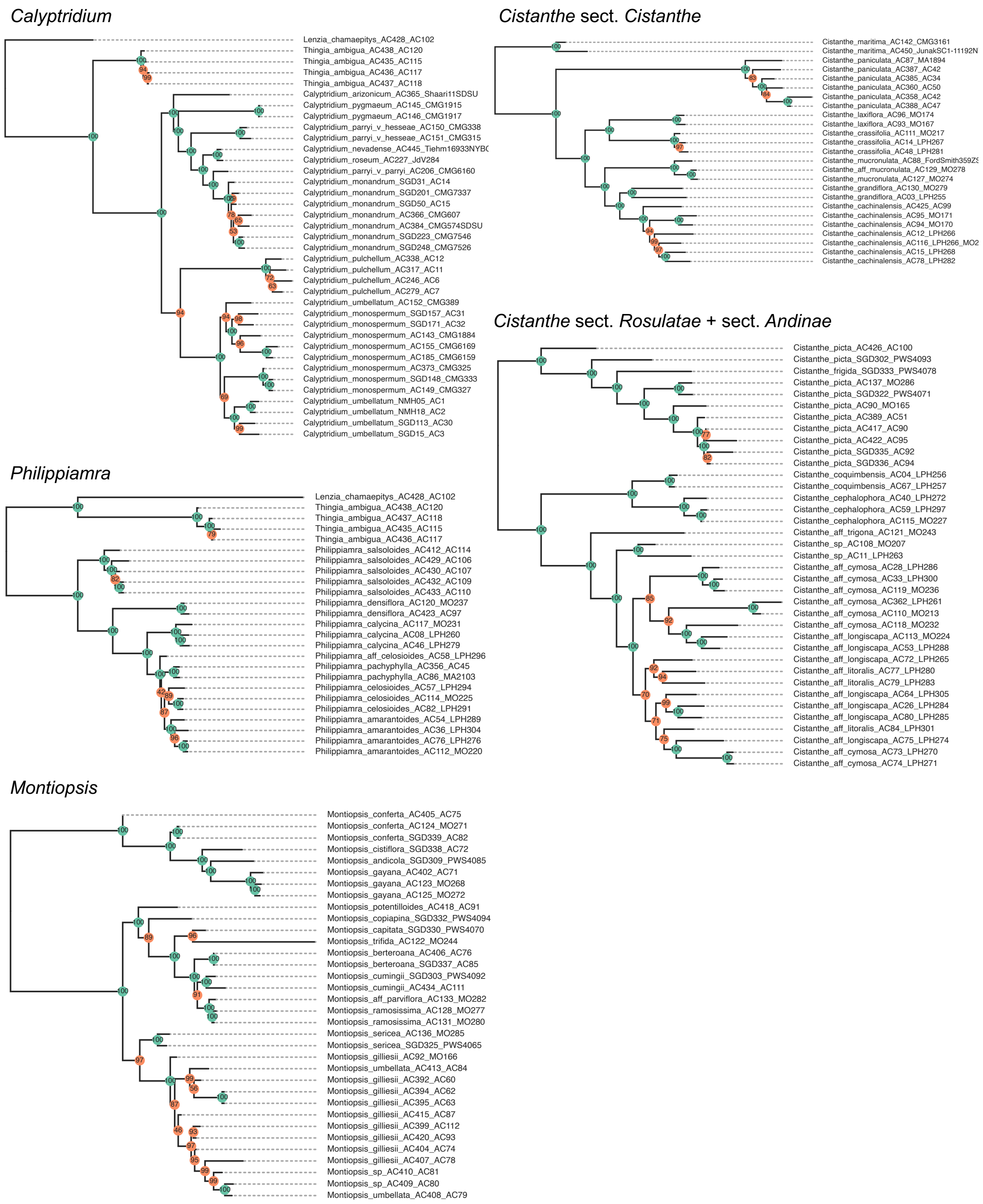


**Sup. Fig. 1.** Subclade-level Maximum-Likelihood (ML) phylogenies using coding sequence-referenced ddRAD assemblies. Subclades include: *Calyptridium* (top left), *Philippiamra* (middle left), *Montiopsis* (bottom left left), *Cistanthe* sect. *Cistanthe* (top right), and *Cistanthe* sect. *Rosulatae* + sect. *Andinae* (bottom right). The trees were inferred using IQ-TREE2 with 1,000 Ultrafast Bootstraps, with support values on nodes (support = 100 in green; support < 100 in orange).

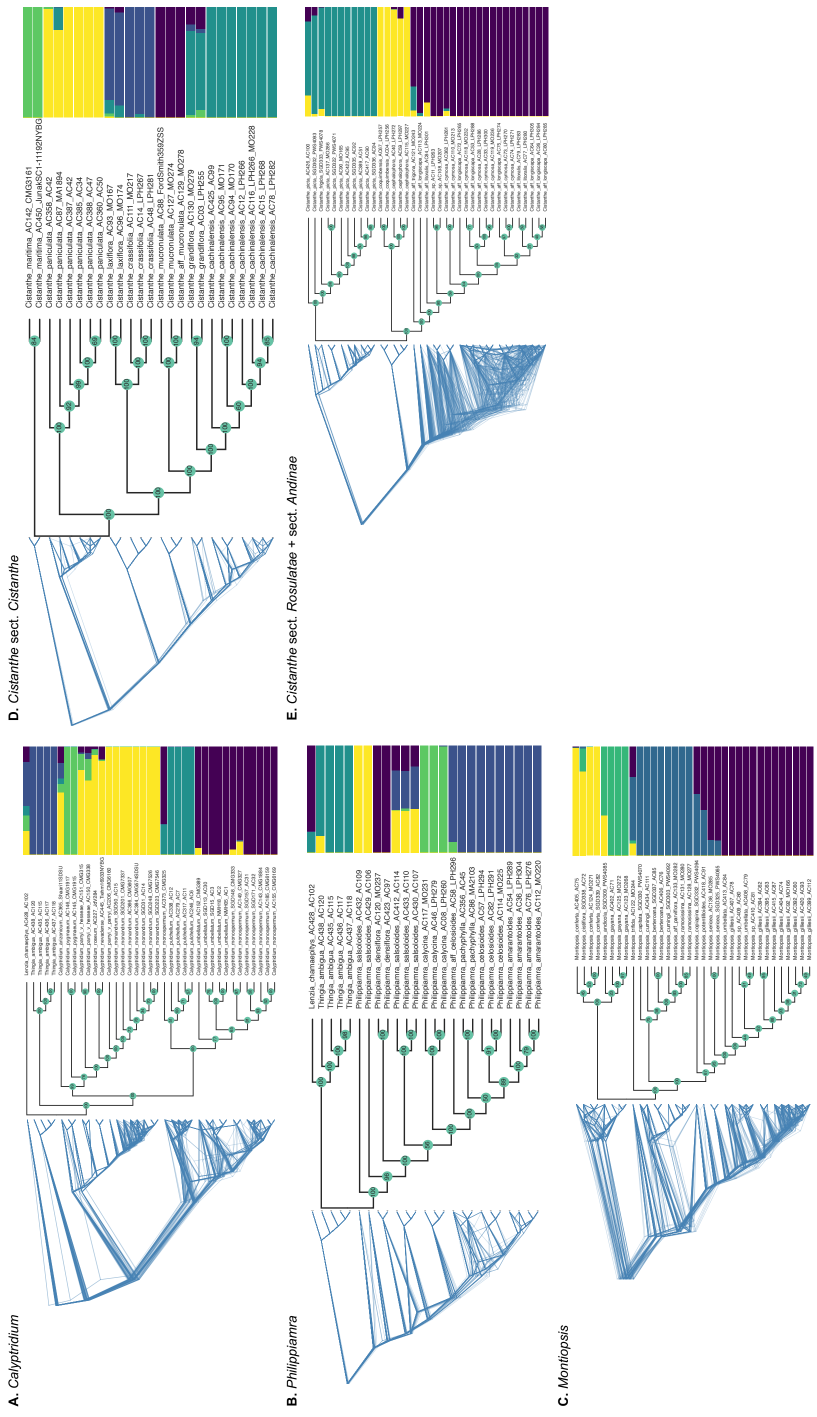


**Sup. Fig. 2.** Subclade-level Tetrad phylogenies (coalescent-based quartet trees) and genomic structure using coding sequence-referenced ddRAD assemblies. **A**. *Calyptridium*, **B**. *Philippiamra*, **C**. *Montiopsis*, **D**. *Cistanthe* sect. *Cistanthe*, and **E**. *Cistanthe* sect. *Rosulatae* + sect. *Andinae*. Leftmost tree in each panel shows a sample of 100 out of 1,000 quartet-based bootstrap replicates, and the middle tree shows the consensus Tetrad phylogeny. The rightmost graph shows genetic clustering, as inferred by ancestry coefficients estimated with sparse nonnegative matrix factorization (sNMF).

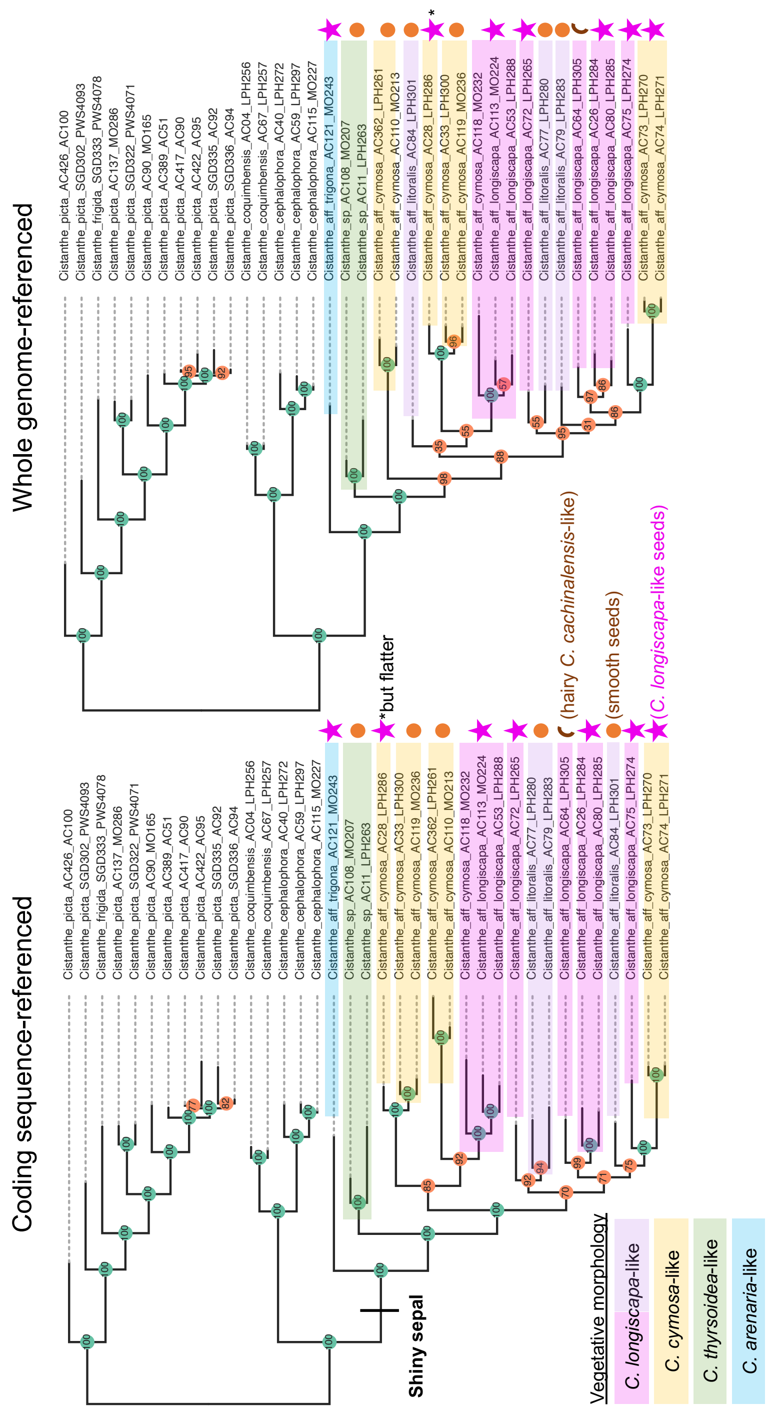


**Sup. Fig. 3.** Distribution of morphological characters in the *Cistanthe longiscapa* complex. Tips in both the coding sequence-referenced and genome-referenced ML trees are highlighted according to the likeness of the specimens' vegetative morphology to what is described in literature as *C. longiscapa*, *C. cymose*, *C. thyrsoidea*, or *C. arenaria*. Tips are additionally annotated with the seed morphology using a pink star symbol (*C. longiscapa*-like pusticulate-tomentose seed), a burnt-orange oval (smooth seeds), or a brown half-circle (hairy seeds).

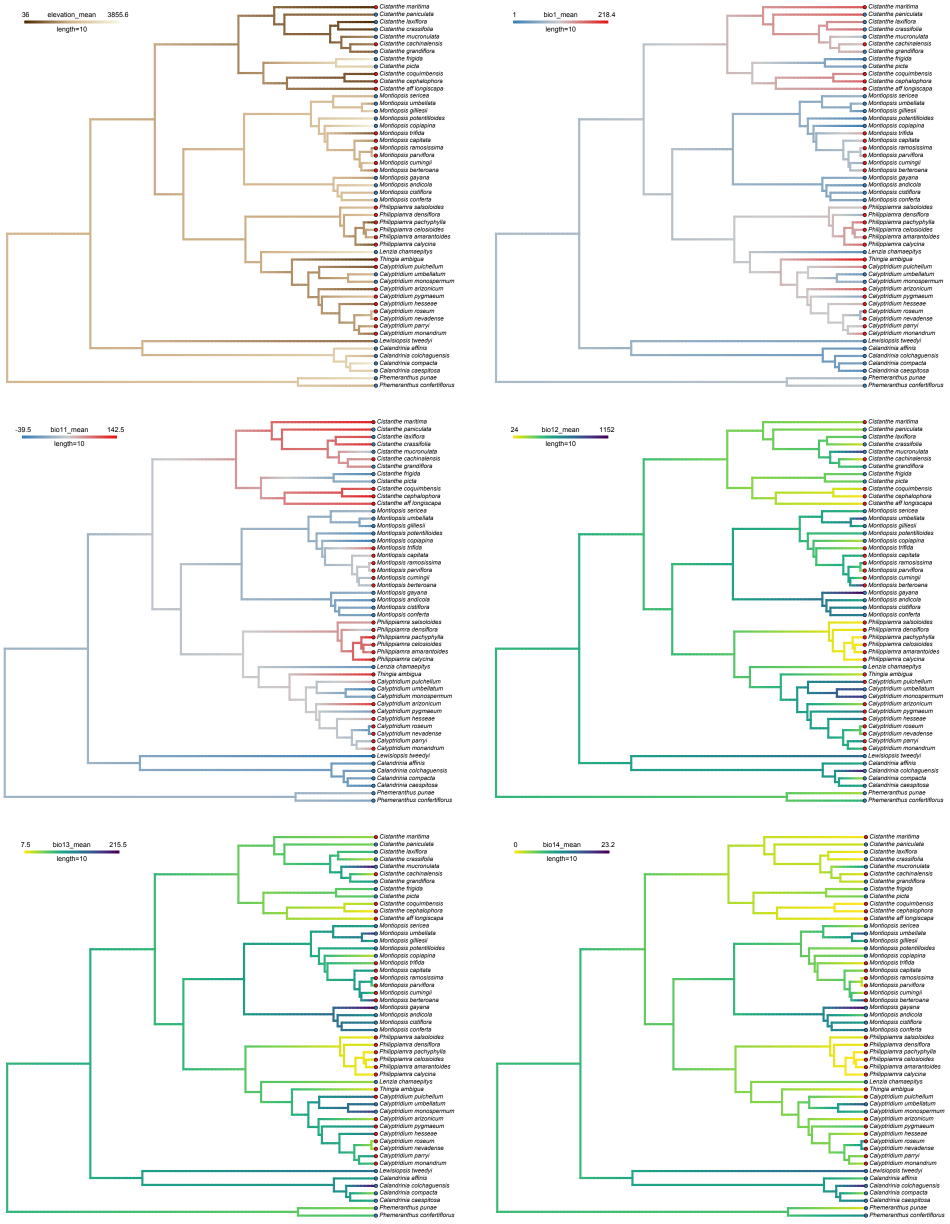


**Sup. Fig. 4.** Ancestral state reconstructions of bioclimatic variables across Cistantheae. Mean species values are used to map elevation (top left), Bio 1 (Mean Annual Temperature; top right) Bio11 (Mean Temperature of Coldest Quarter; middle left), Bio12 (Annual Precipitation; middle right), Bio13 (Precipitation of Wettest Month; bottom left), and Bio14 (Precipitation of the Driest Month) onto the tree with Ornstein-Uhlenbeck (OU) models. Tips are annotated with life history: red circles for annuals and blue circles for perennials. Darker brown ranch colors represent lower mean elevation (m) and lighter colors represent. For Bio1 and Bio11, branch colors represent temperature values (ºC*100), with warmer colors (red) indicating higher temperatures and colder colors (blue) indicating lower temperatures. For the precipitation-related variables, branch colors range from yellow (lower precipitation) to purple (higher precipitation), measured in mm.

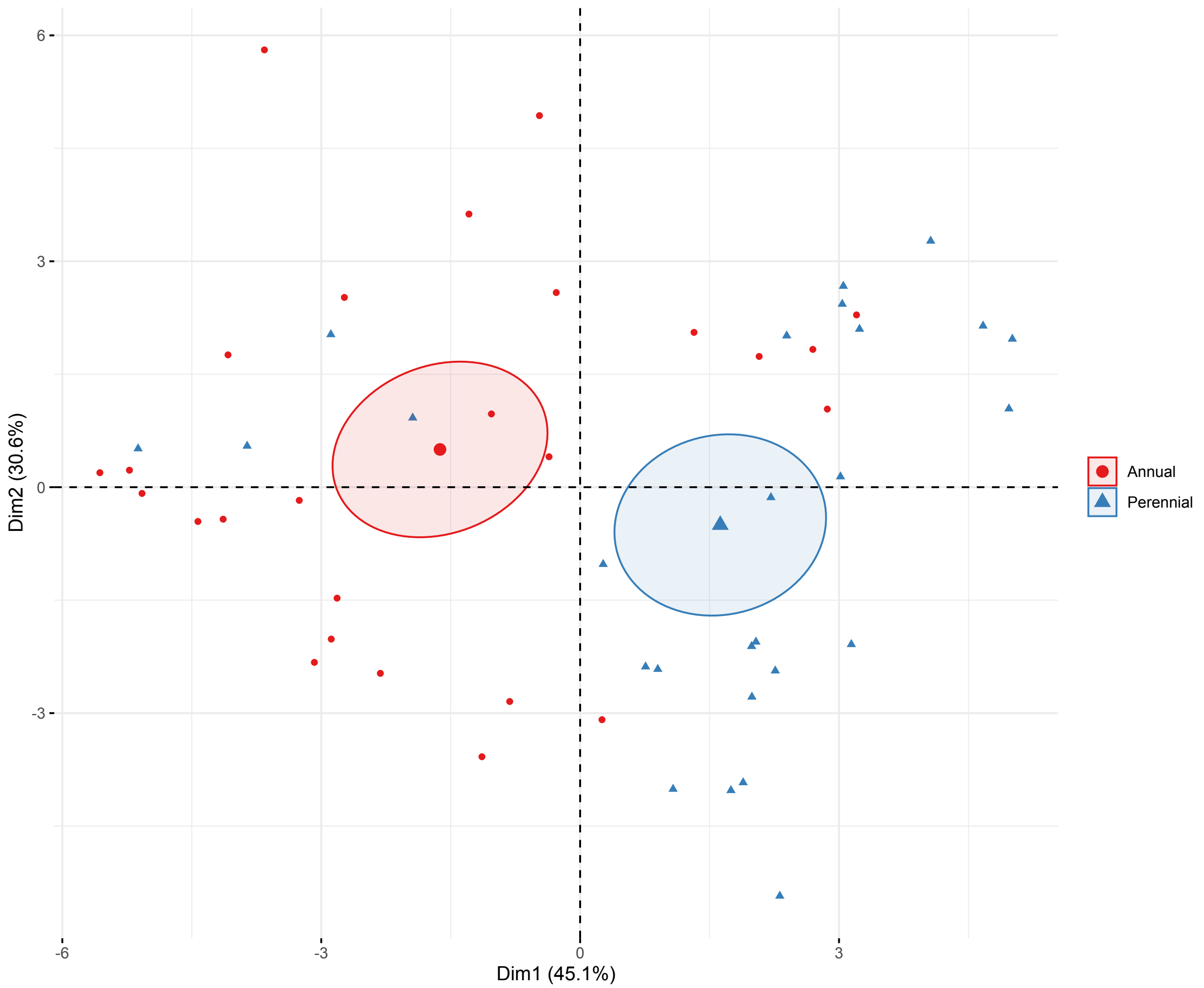


**Sup. Fig. 5.** PCA of bioclimatic variables. Species are plotted along the first two PC axes, which explain 45.1% and 20.6% of the total variance. Annual species are represented in red circles, perennials in blue triangles. 95% confidence ellipses around the annual and perennial species centroids show separation along Dim1, suggesting differentiation between life history strategies.

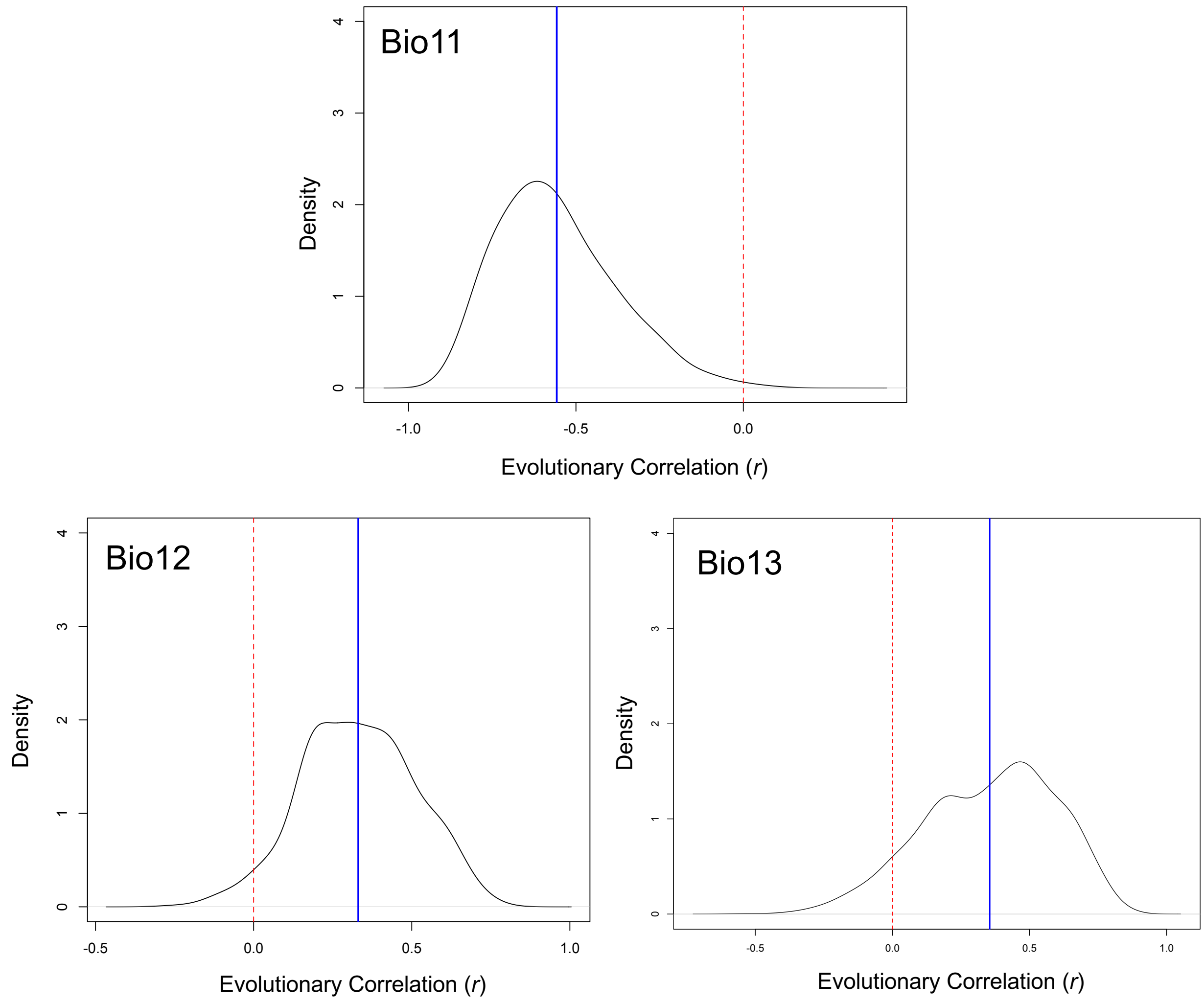


**Sup. Fig. 6.** Posterior density distributions of evolutionary correlations between life history and climate variables across Cistantheae. Vertical red dashed lines denote where evolutionary correlation r = 0; solid blue lines denote the posterior mean. The posterior mean was negative in Bio11 (Mean Annual Temperature; r = -0.560; Effective Sample Size, or ESS, of 252.1; top panel), and positive in Bio12 (Annual Precipitation; r = 0.33; ESS = 216.5; bottom left) and Bio13 (Precipitation of Wettest Month; r = 0.34; ESS = 267.8; bottom right); only the evolutionary correlation in Bio11 is deemed significant based on the 95% credible interval (-0.83 – -0.17).

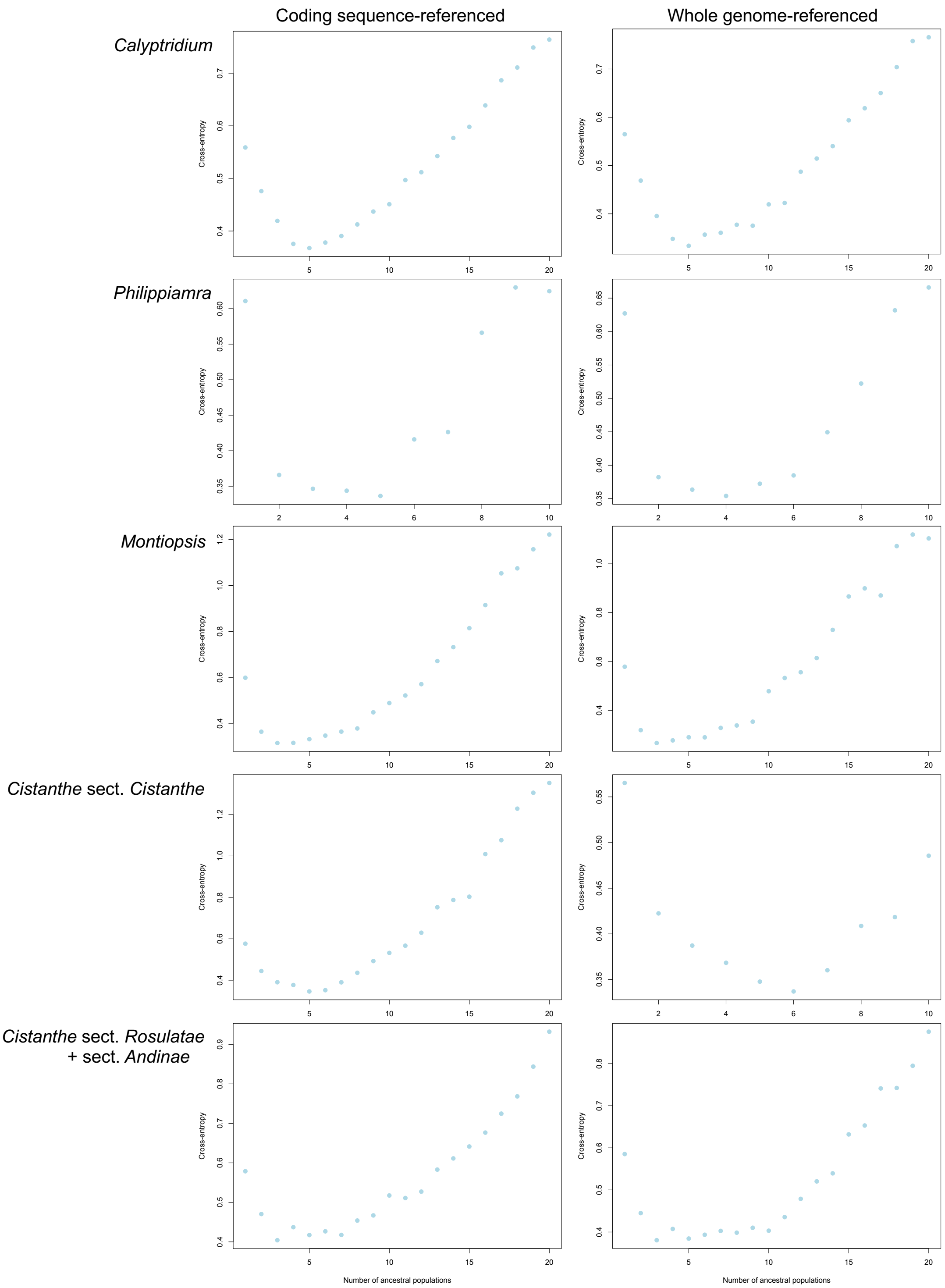


**Sup. Fig. 7.** Cross-entropy estimates from sparse nonnegative matrix factorization (sNMF) analyses for each subclade. Each panel shows the results of sNMF runs for a given subclade, with the number of ancestral populations on the x-axis and the corresponding cross-entropy value on the y-axis. Analyses were conducted using both coding-sequence referenced assemblies (left column) and whole genome-referenced assemblies (right column).

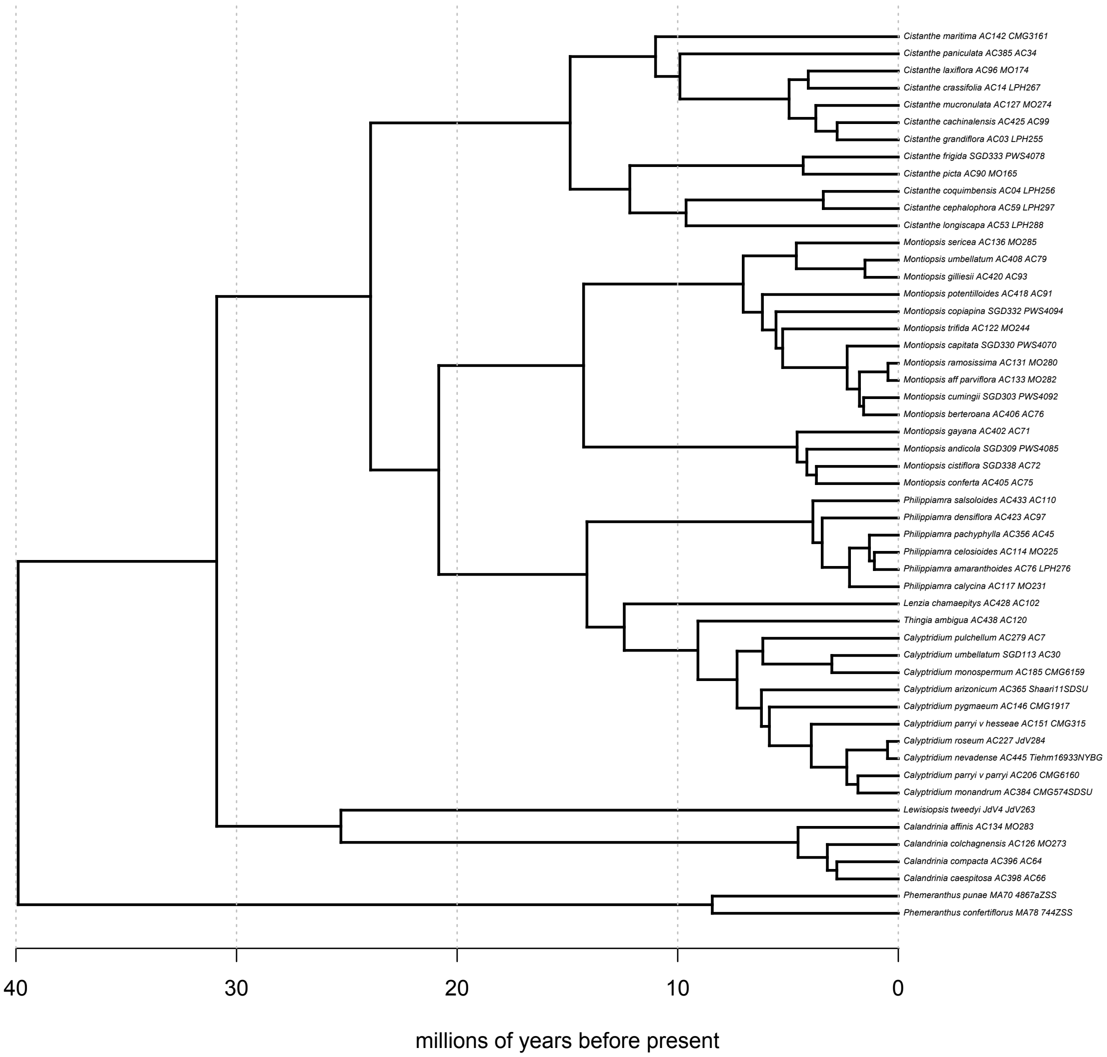


**Sup. Fig. 8.** Time-calibrated phylogeny of representative Cistantheae species. The phylogeny was calibrated using secondary calibrations. Divergence times were estimated under a discrete clock model with 10 rate categories and a smoothing parameter (λ = 0.1). Time is shown in millions of years before present.

**Sup. Table 1.** Summary statistics from phylogenetic generalized least squares (PGLS) and Ornstein-Uhlenbeck phylogenetic linear models (phylolm) between life history and bioclimatic variables.

| Variable | Description | PGLS coefficient | PGLS standard error | PGLS *t* value | PGLS *p* value | PGLS R squared | PGLS lambda | PGLS adjusted *p* value | Phylolm coefficent | Phylolm standard error | Phylolm *t* value | Phylolm *p* value | Phylolm pseudo R squared | Phylolm adjusted *p* value |
|---|---|---|---|---|---|---|---|---|---|---|---|---|---|---|
| Bio1 | Mean Annual Temperature (C*100) | -57.281 | 13.157 | -4.354 | 0.000 | 0.275 | 0.589 | 0.001 | -59.235 | 15.179 | -3.903 | 0.000 | 0.273 | 0.001 |
| Bio2 | Mean Diurnal Range (C*100) | 2.797 | 5.240 | 0.534 | 0.596 | 0.006 | 0.778 | 0.627 | - | - | - | - | - | - |
| Bio3 | Isothermality | -0.675 | 2.010 | -0.336 | 0.738 | 0.002 | 0.853 | 0.738 | - | - | - | - | - | - |
| Bio4 | Temperature Seasonality | 276.055 | 371.049 | 0.744 | 0.460 | 0.011 | 0.978 | 0.512 | - | - | - | - | - | - |
| Bio5 | Max Temp.of Warmest Month (C*100) | -49.160 | 16.136 | -3.047 | 0.004 | 0.157 | 0.795 | 0.007 | - | - | - | - | - | - |
| Bio6 | Min Temp. of Coldest Month (C*100) | -57.875 | 14.778 | -3.916 | 0.000 | 0.235 | 0.654 | 0.002 | - | - | - | - | - | - |
| Bio7 | Temperature Annual Range (C*100) | 10.018 | 13.332 | 0.751 | 0.456 | 0.011 | 0.987 | 0.512 | - | - | - | - | - | - |
| Bio8 | Mean Temp. of Wettest Quarter (C*100) | -51.227 | 14.605 | -3.508 | 0.001 | 0.197 | 0.471 | 0.003 | - | - | - | - | - | - |
| Bio9 | Mean Temp. of Driest Quarter (C*100) | -51.227 | 14.605 | -3.508 | 0.001 | 0.197 | 0.471 | 0.003 | - | - | - | - | - | - |
| Bio10 | Mean Temp. of Warmest Quarter (C*100) | -54.165 | 14.702 | -3.684 | 0.001 | 0.214 | 0.719 | 0.003 | - | - | - | - | - | - |
| Bio11 | Mean Temp. of Coldest Quarter (C*100) | -61.262 | 14.260 | -4.296 | 0.000 | 0.270 | 0.713 | 0.001 | -63.911 | 16.089 | -3.972 | 0.000 | 0.273 | 0.001 |
| Bio12 | Annual Precipitation (mm) | 242.630 | 89.571 | 2.709 | 0.009 | 0.128 | 0.344 | 0.014 | 229.870 | 95.391 | 2.410 | 0.020 | 0.140 | 0.020 |
| Bio13 | Precip. of Wettest Month (mm) | 46.974 | 14.923 | 3.148 | 0.003 | 0.165 | 0.000 | 0.006 | 46.814 | 16.288 | 2.874 | 0.006 | 0.165 | 0.007 |
| Bio14 | Precip. of Driest Month (mm) | 5.319 | 1.678 | 3.170 | 0.003 | 0.167 | 0.203 | 0.006 | 5.691 | 1.692 | 3.363 | 0.001 | 0.191 | 0.003 |
| Bio15 | Precipitation Seasonality (mm) | -14.078 | 6.109 | -2.304 | 0.025 | 0.096 | 0.763 | 0.034 | - | - | - | - | - | - |
| Bio16 | Precip. of Wettest Quarter (mm) | 125.421 | 42.065 | 2.982 | 0.004 | 0.151 | 0.000 | 0.008 | - | - | - | - | - | - |
| Bio17 | Precip. of Driest Quarter (mm) | 19.099 | 6.561 | 2.911 | 0.005 | 0.145 | 0.312 | 0.009 | - | - | - | - | - | - |
| Bio18 | Precip. of Warmest Quarter (mm) | 18.135 | 9.493 | 1.910 | 0.062 | 0.068 | 0.786 | 0.077 | - | - | - | - | - | - |
| Bio19 | Precip. of Coldest Quarter (mm) | 113.426 | 46.504 | 2.439 | 0.018 | 0.106 | 0.296 | 0.026 | - | - | - | - | - | - |
| Elevation | - | 938.846 | 291.143 | 3.225 | 0.002 | 0.172 | 0.417 | 0.006 | 854.391 | 268.061 | 3.187 | 0.002 | 0.166 | 0.004 |
